# Cytochrome P450 26b1-mediated specification of vestibular striola and central zones is required for transient responses in linear acceleration

**DOI:** 10.1101/726232

**Authors:** Kazuya Ono, James Keller, Omar López Ramírez, Antonia González Garrido, Omid Zobeiri, Vanessa Chang, Sarath Vijayakumar, Andrianna Ayiotis, Gregg Duester, Charles C. Della Santina, Sherri M. Jones, Kathleen Cullen, Ruth Anne Eatock, Doris K. Wu

**Affiliations:** National Institute on Deafness and Other Communication Disorders, National Institutes of Health, Bethesda, MD 20892, USA; Department of Neurobiology, University of Chicago, Chicago, IL 60637, USA; Department of Physiology McGill University, Montreal, QC H3G 1Y6, Canada; Department of Special Education and Communication Disorders, 301 Barkley Memorial Center, University of Nebraska-Lincoln, Lincoln, NE 68583-0738, USA; Department of Otolaryngology - Head and Neck Surgery, Johns Hopkins University School of Medicine, Baltimore, MD 21205, USA; Neuroscience and Aging Research Center, Stanford Burnham Prebys Medical Discovery Institutes, CA 92037, USA; Department of Biomedical Engineering, Johns Hopkins University School of Medicine, Baltimore, MD 21205, USA

## Abstract

Each vestibular sensory epithelia of the inner ear is divided into two zones, the striola and extrastriola in maculae of otolith organs and the central and peripheral zones in cristae of semicircular canals, that differ in morphology and physiology. We found that formation of striolar/central zones during embryogenesis requires Cytochrome P450 26b1 (Cyp26b1)-mediated degradation of retinoic acid (RA). In *Cyp26b1* conditional knockout mice, the identities of the striolar/central zones were compromised, including abnormal innervating neurons and otoconia in otolith organs. Vestibular evoked potentials (VsEP) in response to jerk stimuli were largely absent. Vestibulo-ocular reflexes and standard motor performances such as forced swimming were unaffected, but mutants had head tremors and deficits in balance beam tests that were consistent with abnormal vestibular input. Thus, degradation of RA during embryogenesis is required for patterning highly specialized regions of the vestibular sensory epithelia that may provide acute feedback about head motion.

## INTRODUCTION

The sense of balance is mediated by the integration of vestibular, visual, and proprioceptive systems. While unilateral vestibular deficits can largely be compensated by sensorimotor reorganization, bilateral vestibular damage, caused by systemic insults such as aminoglycoside ototoxicity, often cannot^1^. As a consequence, patients with chronic vestibulopathy are disabled by imbalance and oscillopsia. Understanding how vestibular inputs are encoded by vestibular organs to maintain gaze and head stability is important from basic science and therapeutic perspectives.

Vestibular sensory epithelia are comprised of the maculae of the utricle and saccule, which detect linear acceleration, as well as three canal cristae, which detect angular acceleration. These sensory epithelia each have two types of mechano-sensitive hair cells (HCs), type I and type II, which are surrounded by supporting cells (SCs) and innervated by afferent neurons of the vestibular ganglion (Fig. 1a). Type I and II HCs are contacted by different afferent synaptic terminals: large calyceal endings on type I HCs and small bouton endings on type II HCs. The mechanosensitive hair stereociliary bundles of HCs couple to an otolithic membrane in the utricle and saccule and to a cupula in the cristae. Head accelerations deflect these accessory structures and the coupled stereociliary bundles, modulating mechanotransduction channels in the bundles and ultimately changing the firing rate of afferent neurons^2^.

**Fig. 1.**
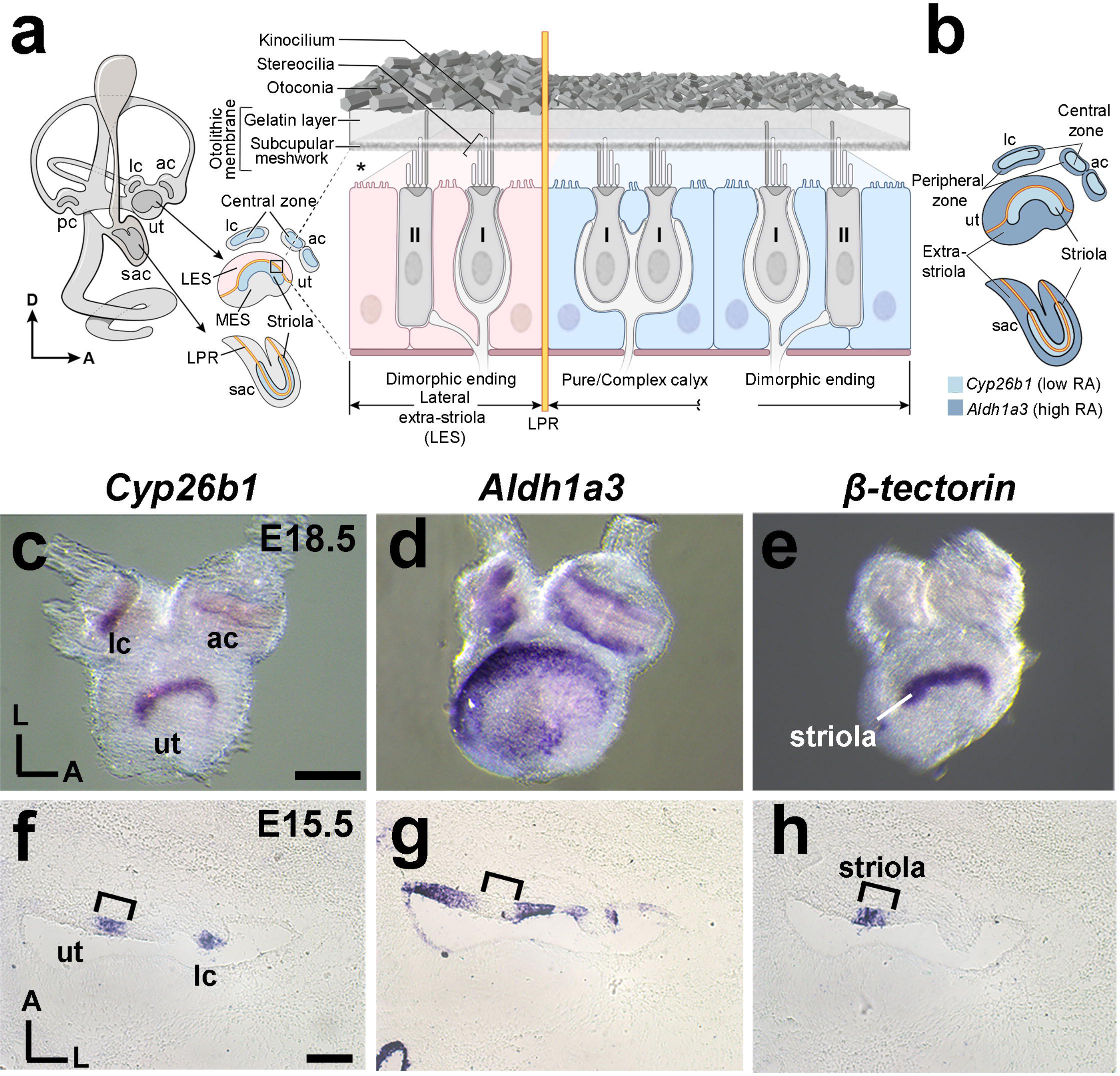
Complementary expression patterns of *Cyp26b1* and *Aldh1a3*. (**a**) Schematic illustration of the inner ear and sectional view of the utricle (ut) across the striola and lateral extrastriola region (LES). Pear-shaped type I HCs are innervated by calyces and cylindrical-shaped type II HCs are innervated by bouton type endings. Pure/complex calyces are exclusively present in the striolar/central zone, whereas dimorphic nerve endings are found across the entire organ. Otoconia are smaller in size and less abundant in striola than extra-striola of the utricle. Asterisk indicates that the relationship between stereocilia bundles and the otoconial membrane is not clear, but evidence suggests that hair bundles of striolar HCs are less firmly embedded in the otolithic membrane than their counterparts in the extra-striolar region^7, 69^. Yellow line represents the line of polarity reversal (LPR), which separates each macula into two regions, LES and MES (medial extrastriola), with opposite hair bundle orientation. The striola of maculae and the central zone of anterior (ac) and lateral cristae (lc) are in blue color. The LPR is at the lateral edge of the striola in the ut but it bisects the striola in the saccule (sac). (**b**) Schematic summary of the expession pattern of *Cyp26b1* in the striolar/central zones (light blue) and *Aldh1a3* in the extrastriolar/peripheral zones (dark blue) of vestibular organs described in (**c-h**). (**c**-**e**) Whole mount *in situ* hybridization analysis of *Cyp26b1*, *Aldh1a3*, and *β-tectorin* transcripts at E18.5 mouse ut and ac and lc. Expression of *Cyp26b1* (**c**) is restricted to the central zone of the two cristae and striolar region of the ut that is *β-tectorin*-positive (**e**), whereas *Aldh1a3* (**d**) is predominantly expressed in the peripheral regions. (**f**-**h**) Adjacent tissue sections at the levels of ut and lc at E15.5. (**f**) *Cyp26b1* expression is concentrated in the supporting cell (SC) layer of the central zone of the lc and striola of the ut, comparable to the *β-tectorin* domain (**h**, bracket). (**g**) *Aldh1a3* expression is complementary to that of *Cyp26b1* (**f**) in each organ. Scale bars for both whole mount and section images equal 200 μm. RA, retinoic acid; D, dorsal; A, anterior; L, lateral; pc, posterior crista.

Each vestibular organ has near its center a conserved, specialized region called the striola in the maculae and the central zone in the cristae^3–5^. Striolas and central zones differ from extrastriolas and peripheral zones in many features including stereociliary bundle morphology, ion channel expression and otoconia size^6–8^. Additionally, afferents form complex calyces around multiple type I HCs in greater proportion in striolar/central zones (Fig. 1a)^4, 5, 9, 10^. Such differences give rise to afferent nerve populations with very different spontaneous and evoked physiological responses. Striolar/central zone afferents have more irregular spike timing and more sensitive to higher-frequency head motion than extrastriolar/peripheral afferents^11–13^. Higher densities of low-voltage-activated K (K_LV_) channels are expressed in striolar/central zone afferents, making them less excitable - less likely to fire in response to small currents – and more directly driven by synaptic inputs, which contributes to their irregular firing patterns^14^. By virtue of their different regularities, afferents from the two zones encode head motion into spike trains by different strategies: temporal pattern of spikes for the striolar/central zones vs. spike rate for extrastriolar/peripheral zones^15^, which are optimal for different kinds of sensory information. The distinct anatomy and physiology of the vestibular epithelia zones motivated our interest in understanding how the zones are established during development and their distinct contributions to key vestibular-driven functions such as the vestibulo-ocular and vestibulo-spinal reflexes (VOR and VSR).

We evaluated the importance of the striola to the VsEP evoked by linear jerk stimulation, a method that allows *in vivo* assessment of macular function independently of other sensory and motor contributions^16–18^. Several lines of indirect evidence implicate irregular afferents, which innervate the striola, in generating the VsEP^2, 16, 19, 20^. Here we directly tested this notion by eliminating striolar specializations. We show that specification of the striolar/central zones of vestibular organs requires degradation of retinoic acid (RA) by the zone-specific expression of *Cyp26b1,* a gene encoding a RA degradation enzyme. RA is the bioactive form of vitamin A (retinol) and it serves as a transcription factor that regulates many cellular events during embryonic development^21^. The availability of RA during embryogenesis is controlled by restricted expression of RA synthesizing enzymes such as class 1A aldehyde dehydrogenases (Aldh1a), as well as degradation enzymes such as Cyp26s^21^. The complementary expression patterns of these enzymes during embryogenesis are important in patterning many tissues including the anterior-posterior axis of the inner ear^22–25^. We show that *Cyp26b1* conditional knockout mice (cKO) exhibit a severe reduction of striolar/central zones as manifested by a number of morphological, molecular and physiological properties. These mice have normal horizontal angular vestibular ocular reflexes (aVOR), derived from the horizontal cristae, and normal responses to off-vertical axis rotation (OVAR), derived from the macular organs, but they lack VsEP. They appear normal in their daily activities but display a characteristic head tremor and have difficulty traversing a balance beam. Together, our results suggest that the striola and central zones are not essential for mediating vestibular ocular reflexes. Rather, striola and central zones are important for responding to changes in linear acceleration and are likely important for controlling head stability and performing challenging vestibulo-motor activities.

## RESULTS

### Complementary expression patterns of *Cyp26b1* and *Aldh1a3* in the vestibular sensory organs

We observed complementary expression patterns of transcripts for the RA degrading enzyme Cyp26b1 and the RA synthesizing enzyme Aldh1a3 in developing vestibular organs that were not described previously. During embryogenesis, the sensory epithelia of the utricle (Fig. 1b-d), saccule (Fig. 1b, Supplemental Fig.1) and cristae (Fig. 1b-d) show *Cyp26b1* expression in the center surrounded by *Aldh1a3* expression in the periphery. The *Cyp26b1* expression domain in the maculae appears comparable to that of *β-tectorin*, a striolar SC marker (Fig. 1c,e)^26^. Adjacent cryo-sections processed for *in situ* hybridization confirmed the complementary relationships among *Aldh1a3*, *Cyp26b1* and *β-tectorin* in both maculae and in the lateral crista (Fig.1f-h, Supplemental Fig.1). Expression of all three genes is concentrated in the basal epithelium corresponding to the layer of SC. Together, these expression patterns suggest that differential expression of RA could establish the central and peripheral zones of vestibular organs.

### Specification of the striolar/central zone in vestibular organs requires *Cyp26b1*

To test the hypothesis that decreased RA levels specify the striolar/central zone of each vestibular sensory organ, we analyzed vestibular organs in gain- and loss-of RA function mutants, *Cyp26b1* and *Aldh1a3* knockout mice respectively. The Ca^2+^-binding protein oncomodulin (Ocm) is expressed only in the type I HCs located within the striolar/central zone (Fig. 2a)^27^. In *Cyp26b1^-/-^* inner ears, Ocm expression is reduced in both maculae and cristae (Fig. 2b, Supplemental Fig. 2). Quantification of HC number showed a significant decrease in the percentages of Ocm^+^ HCs in *Cyp26b1^-/-^* utricles (Fig. 2d), saccules (Supplemental Fig. 2) and lateral cristae (Fig. 2e), without apparent loss in the total number of Myo7a-positive HCs in each sensory organ (Fig. 2b).

**Fig. 2.**
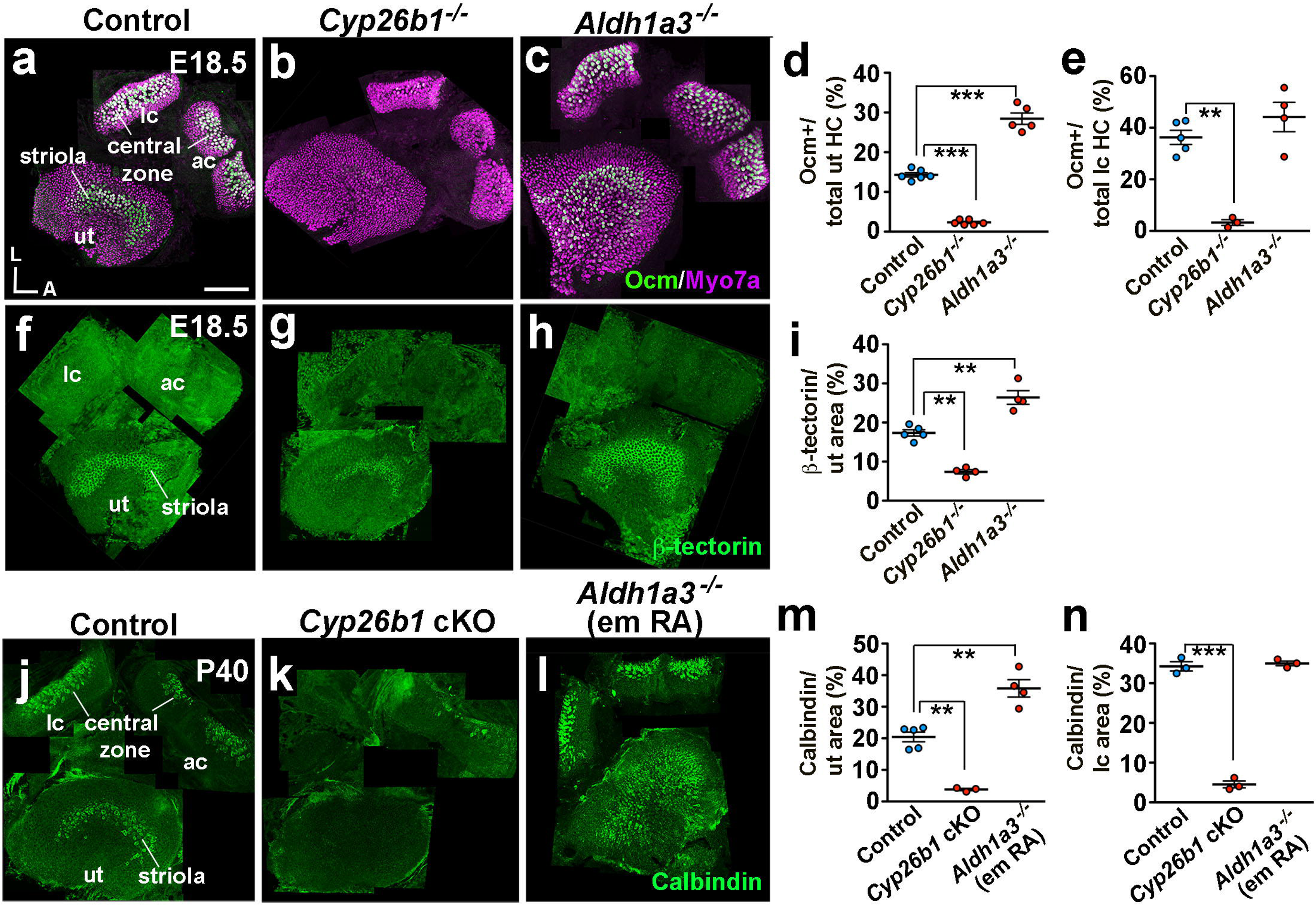
Disruption of RA signaling alters striola and central zone formation in the vestibular sensory organs. (**a**-**c**) Immunostaining using antibodies against myosin7a (magenta) and oncomodulin (Ocm; green) on HCs of E18.5 utricle (ut), anterior crista (ac), and lateral crista (lc). (**a**) In *Cyp26b1^+/-^* or *Aldh1a3^+/-^* controls, Ocm is expressed in type I HCs in striola and central zones only, whereas all HCs are positive for Myosin7a. (**b**-**e**) Ocm expression is severely reduced in *Cyp26b1^-/-^* utricles (**b**, **d**, 2.3 ± 0.2 %, n = 6, P < 0.0001) but expanded in utricles of *Aldh1a3^-/-^* mutants (**c**, **d**, 28.4 ± 1.4 %, n = 5, P < 0.0001) when compared to controls (14.3 ± 0.5 % in controls, n = 6). Ocm^+^ HCs in lateral cristae are reduced in *Cyp26b1^-/-^* mutants (**b**, **e**, 3.2 ± 1.1, n = 3, P = 0.0005) but remain unchanged in *Aldh1a3^-/-^* mutants (**c**, **e**, 44.1 ± 5.6 %, n = 4, P = 0.302), compared to controls (36.2 ± 2.7 %, n = 5). (**f**) Immunostaining of β-tectorin that labels striolar supporting cells in control utricles only. (**g**-**i**) β-tectorin-positive domain is reduced in *Cyp26b1^-/-^* (**g**, **i**, 7.4 ± 0.6 %, n = 4, P = 0.0002) but increased in *Aldh1a3^-/-^* (**h**, **i**, 26.4 ± 1.7 %, n = 4, P = 0.0003) mutant utricles, compared to controls (17.4 ± 0.9 % in controls, n = 5). (**j**-**l**) Immunostaining of afferent nerve endings in the striola and central zones with anti-calbindin antibodies. (**k**-**n**) Calbindin immunoreactivity is severely reduced in *Cyp26b1* cKO (**k**, **m**, 3.8 ± 0.3 %, n = 3, P = 0.0005), but increased in *Aldh1a3^-/-^* (em RA) (**l**, **m**, 36.0 ± 6.6 %, n = 4, P = 0.0006) mutant utricles, compared to controls (**j**, **m**, 19.7 ± 1.7 %, n = 5). Similar to Ocm-positive region, calbindin-positive area is decreased in *Cyp26b1* cKO (**k**, **n**, 4.7 ± 1.4 %, n = 3, P < 0.0001), but unchanged in cristae of *Aldh1a3^-/-^* (em RA) mutant ears (**l**, **n**, 34.4 ± 0.5, n = 3, P = 0.3021), compared to controls (**j**, **n**, 34.2 ± 1.1 n = 3). The one-way ANOVA with multiple comparisons was applied. A, anterior; L, lateral. \*\**P* < 0.01 and \*\*\**P* < 0.001. Scale bar; 200 mm.

In *Aldh1a3^-/-^* mutants, in which RA synthesis is reduced, the Ocm expression domain is increased in the maculae but not cristae (Fig. 2c, Supplemental Fig. 2). In *Aldh1a3^-/-^* utricles, the expression domain of Ocm expands towards the medial extrastriolar (MES) region (Fig.1a) but not the lateral extrastriolar (LES) region of utricles (Fig. 2c, Supplemental Fig. 3); in *Aldh1a3^-/-^* saccules, in contrast, the Ocm expression domain is broadly expanded (Supplemental Fig. 3). The percentages of Ocm^+^ HCs are increased per utricle (Fig 2d) and saccule (Supplemental Fig. 2), compared to controls. The increased Ocm^+^ HC numbers in *Aldh1a3^-/-^* ears are consistent with the decreased numbers in *Cyp26b1^-/-^* ears. In contrast, no increase in Ocm^+^ HCs is evident for the lateral crista (Fig. 2e), suggesting the presence of other compensatory sources of RA in the cristae. Nevertheless, these combined results (Table 1) suggest that formation of the striolar/central zone-specific HCs in vestibular organs is regulated by Cyp26b1-mediated degradation of RA.

**Table 1:**
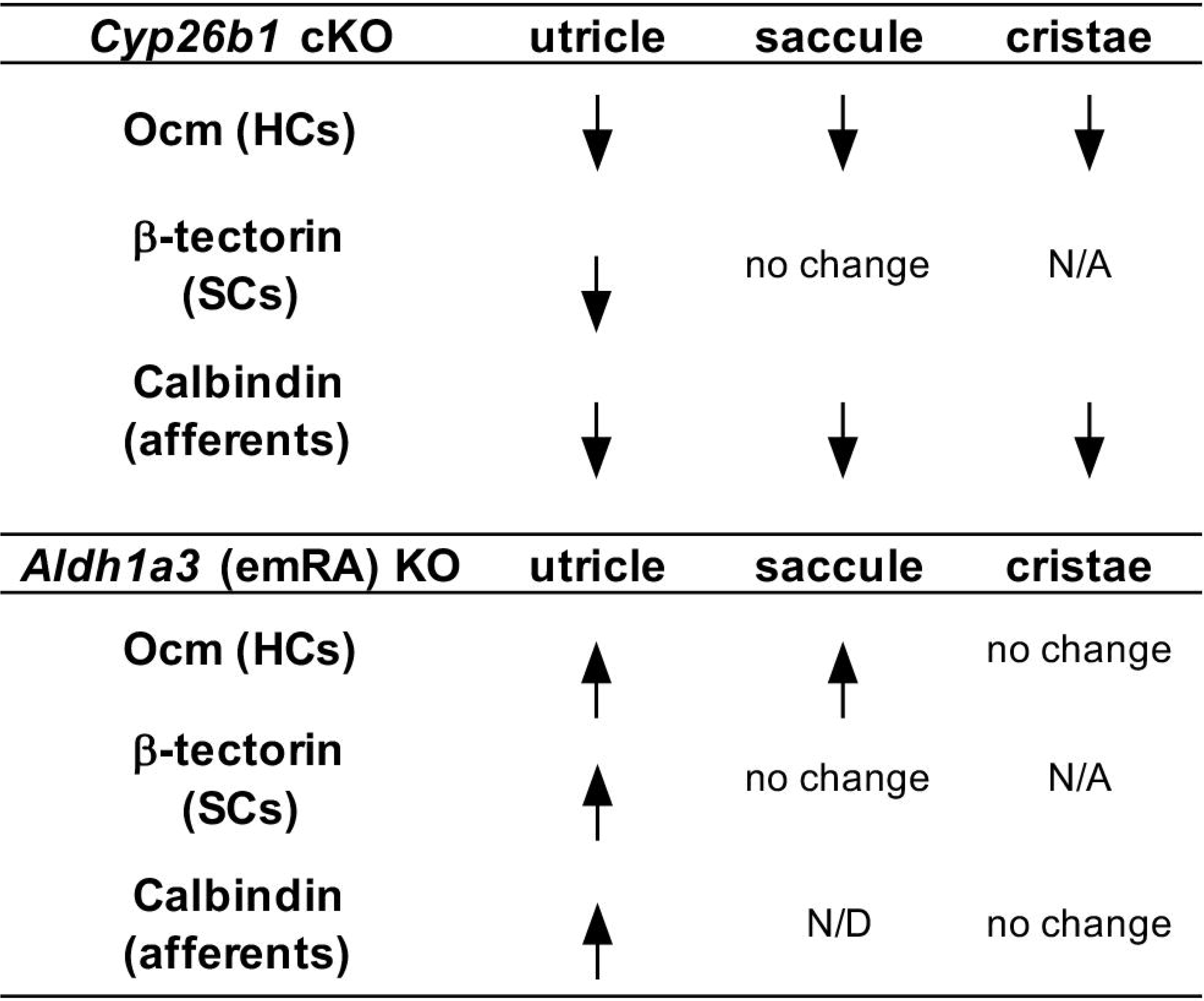
Phenotypic summary of Cyp26b1 cKO and Aldh1a3 (emRA) KO.

| *Cyp26b1* cKO | utricle | sacculle | cristae |
| --- | --- | --- | --- |
| Ocm (HCs) | ↓ | ↓ | ↓ |
| β-tectorin (SCs) | ↓ | no change | N/A |
| Calbindin (afferents) | ↓ | ↓ | ↓ |

| *Aldh1a3* (emRA) KO | utricle | sacculle | cristae |
| --- | --- | --- | --- |
| Ocm (HCs) | ↑ | ↑ | no change |
| β-tectorin (SCs) | ↑ | no change | N/A |
| Calbindin (afferents) | ↑ | N/D | no change |

We next examined whether the identity of SCs in the striolar/central zone of vestibular organs is also affected in mutants deficient for enzymes in the RA pathway. In normal maculae, β-tectorin is specifically expressed in striolar SCs (Fig. 2f, i)^26^. Consistent with the changes in Ocm expression by HCs, the extent of β-tectorin expression is reduced in *Cyp26b1^-/-^* utricles (Fig. 2g, i) and increased in *Aldh1a3^-/-^* utricles (Fig. 2h, i). These results show that striolar SC identity in the utricle requires the decrease in striolar RA levels that is mediated by Cyp26b1. In the saccule, in contrast, no significant difference in the β-tectorin positive-region was detected in either *Cyp26b1^-/-^* or *Aldh1a3^-/-^* ears (Table 1, Supplemental Fig. 2), suggesting that changes in RA levels may not affect SC formation in the saccule. In cristae, which have different accessory structures than the maculae, β-tectorin is not expressed (Fig. 2f) and there is no known SC marker for the central zone, preventing the evaluation of RA effects on SC identity.

Another prominent feature of mammalian vestibular epithelia is the postnatal development of afferent nerve endings (calyces) encasing single or multiple type I HC bodies (Fig. 1a)^6, 9, 10^. In order to investigate the role of RA in calyx formation, we bypassed the lethality of *Cyp26b1^-/-^* and *Aldh1a3^-/-^* mice^28, 29^ by first generating cKO of *Cyp26b1* (*Foxg1^Cre^; Cyp26b1^lox/-^*) using *Foxg1^Cre^,* which is expressed in tissues such as the otic placode and forebrain during early embryogenesis^30^. We also generated viable *Aldh1a3^-/-^* mice by supplementing maternal <u>RA</u> during <u>Em</u>bryonic day(E) 8.5 to E14.5 of development (*Aldh1a3^-/-^* (em RA)). This RA supplementation has been demonstrated to rescue the RA requirements of *Aldh1a3^-/-^* mice during early embryogenesis without reducing the inner ear defects, which develop later^29, 31^. In *Cyp26b1* cKO mice, Ocm and β-tectorin phenotypes are similar to those of the *Cyp26b1* knockouts (data not shown). We used calbindin-D28K immunoreactivity to selectively stain for striolar/central zone afferents^32^ in control and mis-regulated RA mutants at post-natal day (P) 40 or P45. Similar to what was observed for HCs and SCs, the calbindin expression domain is decreased in all *Cyp26b1* cKO vestibular sensory organs compared to controls (Fig. 2j, k, m, n, Supplemental Fig. 2). These results indicate that the development of calyceal-bearing afferents of striolar/central zones requires relatively lower levels of RA signaling regulated by *Cyp26b1*.

In *Aldh1a3^-/-^* (em RA) mice, the calbindin-expressing domain is increased in the utricle but not in the lateral crista (Fig. 2j, l, m, n, Table 1), consistent with the expanded expanded striolar region in the *Aldh1a3^-/-^* mutants. Together, these results suggest that formation of the striolar/central zone in the vestibular organs requires Cyp26b1 enzyme to degrade the RA emanating from the surrounding sensory tissue, which is largely generated by Aldh1a3 in the maculae, though not in the cristae (Table 1).

### Anatomical features of the striola are affected by *Cyp26b1* cKO mice during embryogenesis

Next, we investigated the mature anatomy of the striola in *Cyp26b1* cKO utricles as a result of altered RA signaling during embryogenesis. A significant characteristic of the maculae is the presense of the otoconia, mineralized calcium carbonate crystals embedded above the otolithic membrane (Fig. 1a)^7^. Under low magnification, the otoconial layer in the utricle is more transparent in the striola than the rest of the organ due to the smaller size and number of otoconia (Fig. 1a, 3a). This clear zone is absent in *Cyp26b1* cKO utricles (Fig. 3b). Scanning electron micrographs (SEM) of control utricles show smaller otoconia and more extensive perforations of the underlying meshwork otoconia in the striola (Fig. 3c-c”)^7^ compared to the MES (Fig. 3c’’’). In *Cyp26b1* cKO utricles, SEM revealed no regional differences in otoconial crystals (Fig. 3d-d’’), consistent with low-power brightfield images (Fig. 3b). Crystals throughout *Cyp26b1* cKO utricles were of similar size to control MES crystals (Fig. 3e) indicating that the zonal difference in otoconial size was absent in the *Cyp26b1* cKO utricles.

**Fig. 3.**
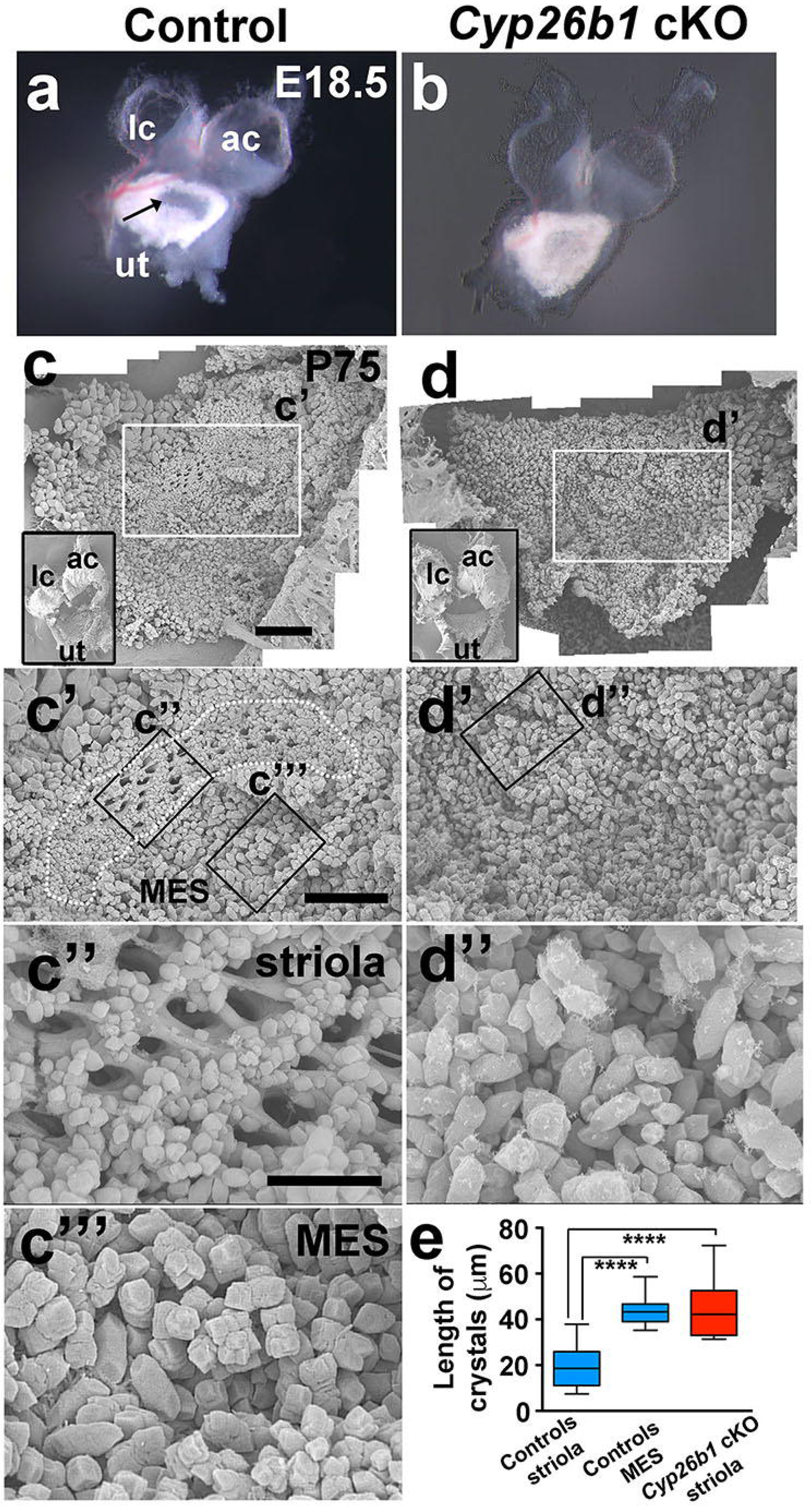
Loss of regional differences in the otoconia of *Cyp26b1* cKO utricles. (**a**, **b**) A dissected utricle (ut), anterior (ac) and lateral cristae (lc) of *Foxg1^Cre^;Cyp26b1^lox/+^* control (**a**) and *Cyp26b1* cKO (**b**) ears at E18.5. In control utricle, the striola region shows a clearance of the otoconia (arrow in **a**), which is missing in *Cyp26b1* cKO utricles. (**c**-**e**) SEM of otoconia in *Foxg1^Cre^;Cyp26b1^lox/+^* controls (**c**-**c”’**) and *Cyp26b1* cKO (**d**-**d”**) utricles at P75. Insets show low power views of respective utricles. In controls (**c’**, **c”**, **c’”**), the striolar region (dotted outline) shows otoconia with smaller crystals (**c”**, **e**, 19.7 ± 1.5 μm, n = 22 crystals) and perforated holes in the subcupular meshwork layer (**c”**), whereas the medial extrastriolar region (MES) shows larger otoconia (**c”’**, **e**, 40.2 ± 2.2 μm, n = 23, P < 0.0001). There is no clear regional difference in the size of the otoconia in the *Cyp26b1* cKO mutant utricle (**d**, **d’**), and the otoconia crystals in the presumptive striolar region are larger in size (**d’’**, **e**, 42.9 ± 3.1 μm, n = 21), comparable to those found in MES of controls (**c”’**, P = 0.6509). The one-way ANOVA with multiple comparisons was applied. \*\*\**P* < 0.001. Scale bars; 200 mm for (**a**, **b**), 300 mm for (**c**, **d**) and (**c’**, **d’**), and 100 mm for (**c’’**, **c”’**, **d”**).

At the sensory epithelium level, the striola has a lower density of HCs (number per surface area) than in the rest of the sensory organ^4^, reflecting larger apical cell surfaces. To investigate whether this regional marker is also affected by RA signaling, the density of HCs in the striolar region was compared between controls and *Cyp26b1* cKO mutants. In *Cyp26b1* cKO utricles, which lack normal striolar identifying features, we delineated a comparable region to the control striola (see Materials and Methods) for comparison. In control utricles, HC density was lower in striola than in LES (Supplemental Fig. 4). This regional difference in HC density was not detected in mutant utricles (Supplemental Fig. 4). Striolar HC density in *Cyp26b1* cKO mutant epithelia was significantly higher than in control striola and comparable to HC density in control LES. Consistently, the ratio of HC number in striola to LES was greater in *Cyp26b1* cKO mutants than in controls (Supplemental Fig. 4), indicating that *Cyp26b1* cKO utricles had more HCs in the striolar region and lacked a zonal difference in HC density.

In control utricles, the ratio of the length of the kinocilium to that of the tallest stereocilium (K/S ratio) is significantly smaller in the striola than in the extrastriola^8^. We therefore measured whether RA signaling affected the K/S. For this analysis, we referenced hair bundle location to the line of polarity reversal (LPR) of stereociliary bundles, which is closely associated with the striola in macular organs (Fig. 1a), and stereociliary bundle orientation was identified by phalloidin staining of actin-filled stereocilia or anti-spectrin staining of the cuticular plate in which stereocila insert^33^. In the mouse, the utricular striola is largely medial to the LPR whereas the saccular striola straddles the LPR (Fig. 1a, Supplemental Fig. 5)^34^. We determined that the relative position of the LPR was not altered in either the mutant utricle or saccule (Supplemental Fig. 5). In control utricles, K/S ratio is higher in the LES than in the striolar region (Supplemental Fig. 6). In contrast, the K/S ratio in the striolar region of *Cyp26b1* cKO utricles was comparable to those of the control utricular LES (Supplemental Fig. 6). These results suggest that the zonal difference in the K/S ratio is absent in the *Cyp26b1* cKO utricles.

Visual inspection of living utricular epithelia with differential contrast microscopy suggested that complex calyces, a prominent features of the striolar/central zone^4, 5, 8^, were less numerous in *Cyp26b1* cKO utricles. To quantitatively examine this impression, we stained neurons with Tuj1 antibody and counted calyces in control and *Cyp26b1* cKO utricular striolas. Complex calyces encasing two or three type I HCs (double or triple calyces, respectively) were present in significantly higher numbers in control striolas (Fig. 4a, c) than in *Cyp26b1* cKO striolar regions (Fig. 4b, c).

**Fig. 4.**
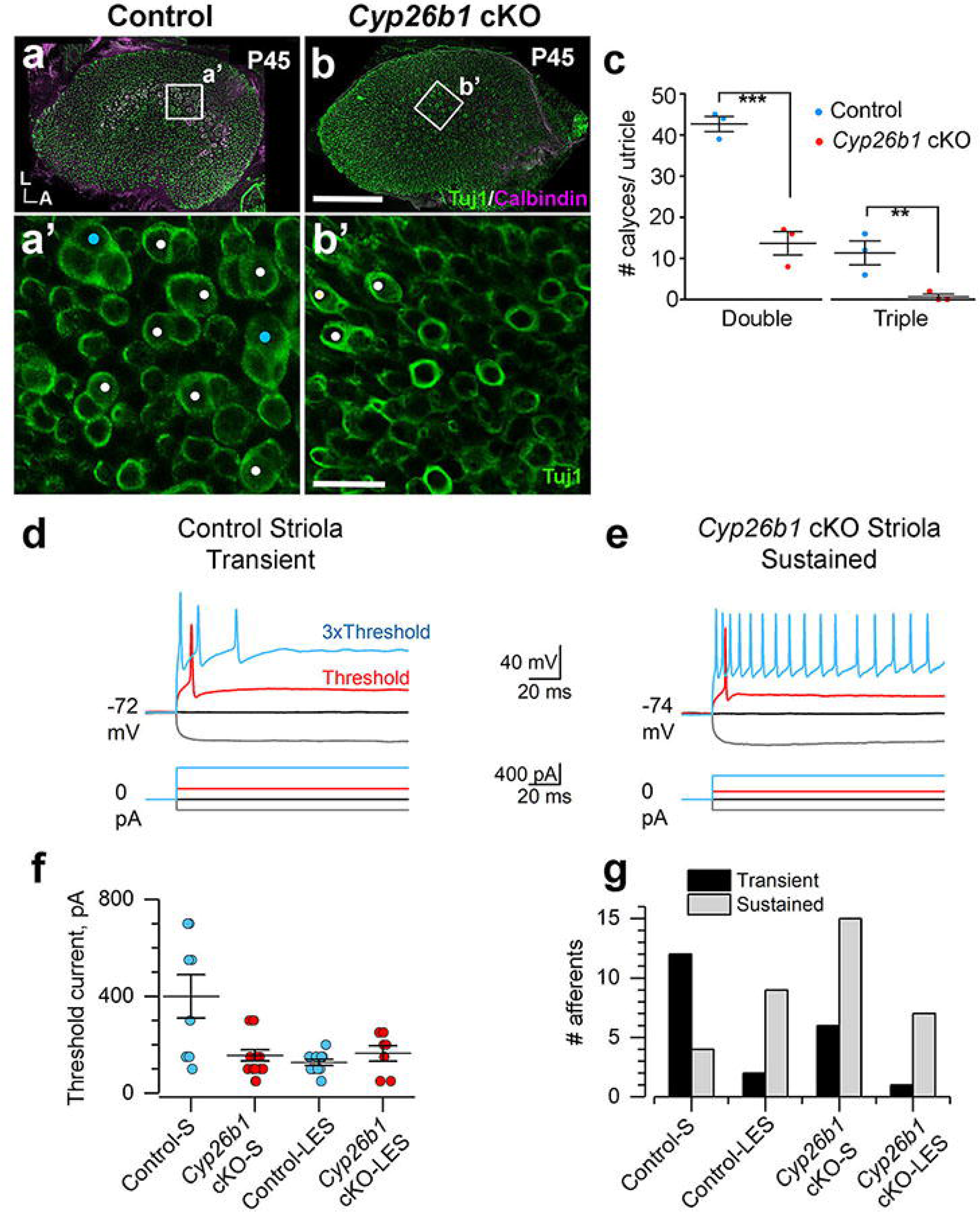
Afferent calyces in *Cyp26b1* cKO utricles showed a reduction in striolar-like morphology and neural activity. (**a**-**c**) P45 wholemount utricles from controls (**a**, **a**’) and *Cyp26b1* cKO (**b**, **b’**) immunolabeled with anti-Tuj1 (green) and anti-calbindin (magenta) antibodies. (**a**, **b**) Maximum intensity projection of the entire utricle. (**a’**, **c**) Enlarged, single plane image of the rectangular region in control striola (**a**), showing the presence of large number of double (white circles, 42.6 ± 1.8 double calyces/ utricle) and triple (cyan circles, 10.0 ± 2.0 triple/utricle, n = 3 utricles) calyces at the cell body level. (**b’**, **c**) Fewer double calyces (white circles, 13.6 ± 2.8 double calyces/utricle, P = 0.001) and no triple calyx (0.6 ± 0.6 triple calyces/utricle, P = 0.006, n = 3) are found in the corresponding region of *Cyp26b1* cKO mutants. Scale bars; 200 μm (**b**); 30 μm (**b’**). Unpaired t-test. **P < 0.01. A, anterior; L, lateral. (**d**-**g**) *Cyp26b1* cKO striolar afferents, like control ES afferents, were more excitable than control striolar afferents, as shown by their tendency to fire more spikes and by reduced current threshold for spiking. (**d, e**) Whole-cell current clamp records from patched calyces of a control striolar afferent (**d**) and a *Cyp26b1* cKO striolar afferent (**e**). 500-ms current steps were delivered in 50-pA increments from –200 pA to >1 nA relative to zero holding current; a subset is shown including the response at threshold current (red). Three times (3x) threshold current (blue) evoked transient spiking in a control striolar afferent but sustained spiking in *Cyp26b1* cKO striolar afferent. (**f**) Mean threshold current was significantly higher in Control-S calyces than in all other categories – i.e., the cKO-S calyces resembled cKO-LES calyces and Control-LES calyces. See text for details on statistics. (**g**) Transient firing was more common in Control-S afferents; sustained firing was more common in Control-LES, *Cyp26b1* cKO-S and *Cyp26b1* cKO-LES afferents.

Together, our results showed that the lack of *Cyp26b1* during embryogenesis affects many mature features of the striola including the otoconia, HC density, hair bundle morphology, expression of Ca^2+^-binding proteins, and incidence of complex calyces, suggesting that striolar maturation depends on low levels of RA signaling.

### Loss of functional properties of the striola in *Cyp26b1* cKO utricles

To assess whether zonal differences in physiology were affected by the mutation, we recorded from HCs and afferent calyces in both controls and *Cyp26b1* cKO mice using a semi-intact preparation of the utricle and the distal nerve. We visualized the epithelium from its apical surface with differential-interference-contrast optics, which allow resolution of hair bundles, HC bodies, and calyceal afferent terminals. Because the *Cyp26b1* cKO utricles had lost the zonal markers that we usually use (hair bundle size, HC density as viewed from above, the proportion of calyces that are complex), we relied on the LPR, assigning recordings in the first 10 HC rows medial to the LPR as “striolar”. We performed the whole-cell recording configuration on visually identified type I or type II HCs or on calyceal afferent terminals that surround type I HCs, and measured voltage-gated currents in voltage clamp mode and resting potential and step-evoked voltage changes in current clamp mode. Recorded HCs and calyceal terminals were filled with fluorescent dye (sulforhodamine 101) in the pipette solution, allowing their visualization with fluorescence optics.

In HCs (n = 64, from 21 mice), there were no obvious effects of the *Cyp26b1* cKO manipulation on whole-cell voltage-sensitive conductances, as revealed by currents evoked by voltage steps or voltage responses to current steps. Wildtype type I and type II cells from mice and other amniotes^35^ have outwardly rectifying K^+^ currents with very different voltage dependence: the voltage of half-maximal activation (V_1/2_) is 40-70 mV more negative in type I HCs compared to type II HCs. The unusually negative voltage dependence of the type I-specific K^+^ conductance (g_K,L_) has significant consequences for the size and speed of the receptor potential and appears to be essential for non-quantal transmission at type I-calyx synapses^36^. This key difference was preserved in the striolar zones of *Cyp26b1* cKO utricles: 3-factor ANOVAs on V_1/2_ for HC type (I vs. II), genotype, and epithelial zone yielded a highly significant difference for HC type (F(1,46) = 1348.33, P = 0) but for no other comparison. Thus, the *Cyp26b1* cKO mutation did not appear to affect the electrophysiological differentiation of type I and type II HCs.

Differences were seen, however, when we compared the spiking activity of striolar and LES calyx-bearing afferents in *Cyp26b1* cKO and *Foxg1^Cre^; Cyp26b1^lox/+^* controls. In the LES, calyx-bearing afferents are dimorphic afferents (forming both calyx and bouton terminals, Fig. 1A). In the striola, calyx-bearing afferents are either dimorphic or pure-calyx afferents. We measured the spiking activity of dimorphic (a nerve with both calyx and bouton endings, Fig. 1A) or pure-calyx vestibular afferent neurons via ruptured-patch recordings from their calyceal terminals; spikes initiate on the afferent neurite below the base of the calyx^6^. Striolar afferent neurons were more excitable in *Cyp26b1* cKO mice compared to *Foxg1^Cre^; Cyp26b1^lox/+^* controls, as measured by a lower current threshold for spiking and a tendency to fire a longer train of spikes in response to supra-threshold current steps. Threshold current was defined as the first level at which spiking occurred as applied currents were incremented in 50-pA steps (red traces, Fig. 4d,e). In comparing threshold currents across genotype and zone with 2-factor ANOVA, we saw significant main effects of genotype (control vs. cKO, F(1,41) = 6.1, P = 0.02), zone (striolar vs. extrastriolar, F(1,41) = 10.0, P = 0.003) and their interaction (F(1,41) = 11.2, P = 0.002). The significant differences were all between control striolar afferents (400 ± 90 pA, n = 8) and each of the other groups (Fig. 4f), including control ES afferents (127.3 ± 12 pA, n = 11; Tukey means comparison: p = 0.0003, effect size (Hedge’s g) 1.9) and *Cyp26b1* cKO striolar afferents (156.3 ± 23 pA, n = 16; P = 0.0005, effect size (Hedge’s g) 1.62). Threshold currents did not differ significantly between *Cyp26b1* cKO striolar afferents and control ES afferents (Tukey means comparison, P = 0.94) or cKO ES afferents (164 ± 32 pA, n = 7, Tukey means comparison, P > 0.99). Thus, the *Cyp26b1* cKO manipulation reduced threshold current for spiking in striolar afferents to levels typical for extrastriolar afferents.

Afferent firing patterns were classified for current steps at 2-3 times threshold current as either *transient (*1-3 onset spikes) or *sustained* (>4 spikes). The tendency of isolated neurons to respond to injected current steps with transient or sustained responses correlates with their tendency to fire more irregularly or more regularly *in vivo*, when they are driven by synaptic inputs from HCs^14^. In control utricles, striolar afferents were more likely to have transient than sustained responses and extrastriolar afferents were more likely to have sustained than transient responses (Fig. 4g), consistent with previous data from rat vestibular ganglion neurons^14^ and calyceal terminals^37^. In *Cyp26b1* cKO utricles, the normal zonal difference in firing pattern disappeared: most striolar afferents, like extrastriolar afferents, had sustained responses to current steps (Fig. 4g).

These changes in excitability in the cKO zone corresponding to the striola (reduced current threshold, more sustained firing) likely involves loss of striolar-specific ion channel expression. Candidates include low-voltage-activated K (K_LV_) channels, which reduce neuronal excitability by increasing K^+^ conductance around resting potential and, in normal utricular afferents, are expressed more in striola than extrastriola^6, 14, 37, 38^. Preliminary comparisons suggest that *Cyp26b1* cKO afferents lack this zonal difference: mean I_KLV_ did not differ significantly across zones (striola: 870 ± 114 pA, n = 16; extrastriola:1192 ± 151 pA, n = 6; P = 0.14, n = 22, estimated power 0.3). Thus, *Cyp26b1 cKO* utricles showed changes in afferent physiology that are consistent with a loss of striolar identity.

### Absent vestibular-evoked potential in *Cyp26b1* cKO mice

To assess the functional consequences of loss of the striola *in vivo*, we recorded with scalp electrodes the vestibular-evoked potential (VsEP), which are thought to represent summed far-field afferent responses to transient linear accelerations, in control and *Cyp26b1* cKO animals. Control animals (*Cyp26b1^lox/+^* and *Foxg1^Cre^; Cyp26b1^lox/+^* heterozygous animals) had normal VsEP responses, whereas *Cyp26b1* cKO mice showed either absent or remnant VsEP responses (Fig. 5a). VsEP thresholds between the *Cyp26b1^lox/+^* and *Foxg1^Cre^; Cyp26b1^lox/+^* heterozygotes were comparable (Fig. 5b). There were also no differences in VsEP response activation latencies (P1, N1) or amplitude (P1-N1) at the highest stimulus level (+6 dB re: 1g/ms) across the two genotypes (data not shown). Taken together, these data show that VsEP was severely affected in *Cyp26b1* cKO mice, indicating that the loss of striola effectively abolishes most, if not all, VsEP response, supporting the existing hypothesis^2^ that the VsEP is a summated potential reflecting the activity of striolar afferents and their downstream targets.

**Fig. 5.**
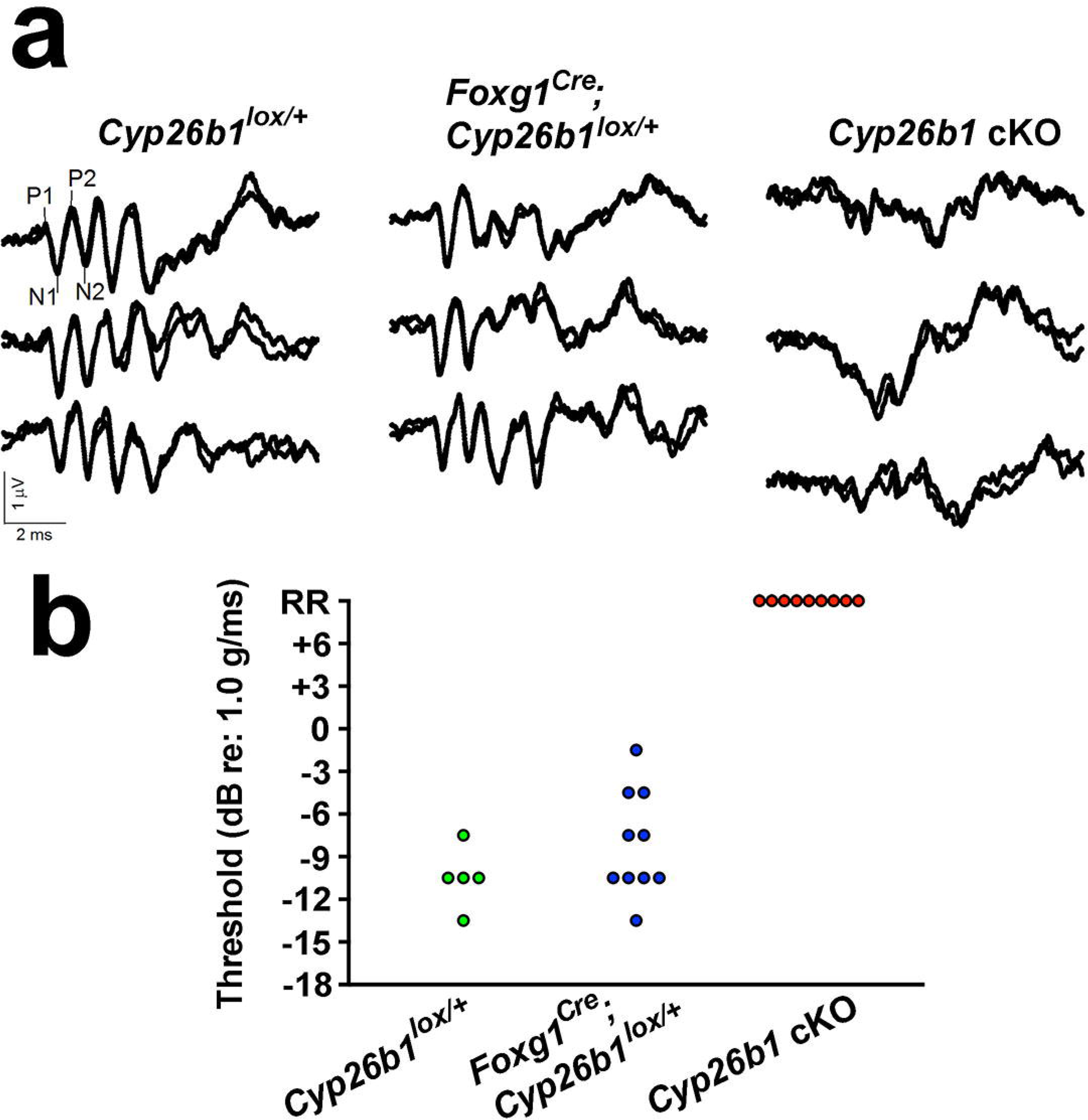
Absence of linear vestibular-evoked potential (VsEP) in *Cyp26b1* cKO mutant mice. (**a**) Three representative VsEP waveforms for each genotype recorded at maximal jerk stimulus (+6 dB). In the *Cyp26b1* cKO mutants, distinct peaks of P1-P2 and N1-N2 are not detectable. (**b**) Summary of thresholds for VsEP determined by various jerk magnitudes. There was no significant difference in VsEP thresholds between the *Cyp26b1^lox/+^* (−10.5 ± 1.34 dB re:1g/ms, n = 5) and *Foxg1^Cre^; Cyp26b1^lox/+^* heterozygotes (−8.1 ± 1.17 dB re:1g/ms; n = 10, P = 0.1026). *Cyp26b1* cKO mutants generated only a remnant response (RR).

### Normal angular vestibulo-ocular reflex and off-vertical axis rotation response in ***Cyp26b1* cKO mice**

Next, we tested whether the VOR was also affected in the *Cyp26b1* cKO mutants based on the hypothesis that the striolar/central zone is important for vestibular-reflex function^13, 20^. The VOR plays an important role in ensuring gaze stabilization during everyday activities by producing compensatory eye movements in the opposite direction of the head movement^39^. Because the dynamic response properties (i.e., gain and phase) of the VOR can be precisely quantified, it has also become a valuable tool for accessing vestibular function in mice. The best characterized VOR is the horizontal angular VOR (aVOR), which counter-rotates eyes in the horizontal plane and is largely driven by signals from the lateral cristae. As a control for eye muscle function, we also examined the optokinetic reflex (OKR), which uses visual signals to control eye motion, allowing tracking of a moving visual scene or objects. Normally, OKR functions in concert with VOR and vestibular neck reflexes to achieve the correct head and eye motion to stabilize visual field^39^. Both the *Foxg1^Cre^;Cyp26b1^lox/+^* heterozygotes and *Cyp26b1* cKO mice showed robust OKR responses (data not shown). Although aVOR gain increased and aVOR phase decreased systematically with increasing frequency of stimulus, there were no significant differences between *Foxg1^Cre^;Cyp26b1^lox/+^* heterozygotes and *Cyp26b1* cKO mice in gain, phase change, model gain or model time constant up to 10 Hz (Fig. 6a-e; P > 0.05 for all comparisons).

**Fig. 6.**
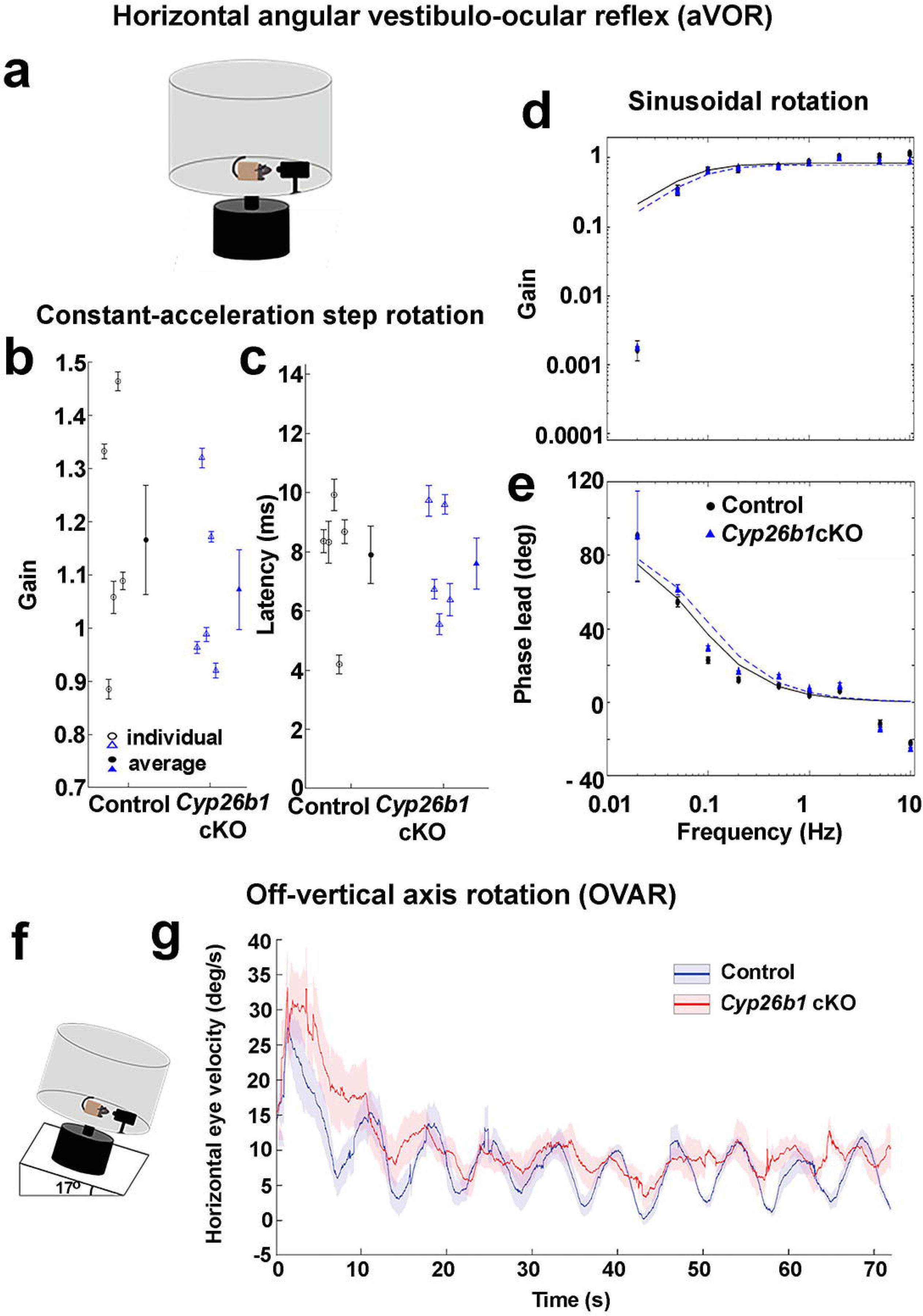
Normal horizontal angular vestibulo-ocular reflex and off-vertical axis rotation response in *Cyp26b1* cKO mice. (**a**) Schematic view of horizontal angular vestibulo-ocular reflex (aVOR) apparatus. Constant-acceleration step gain G_A_ (**b**) and latency (**c**) for aVOR responses to whole-body, 3000°/s^2^ whole-body passive yaw rotations in darkness about an Earth-vertical axis through the head. Open markers denote individual mice; thick markers and lines show mean ± SEM. Gain (**d**) and phase lead (**e**) for yaw slow-phase aVOR responses to 0.02-10 Hz, 100°/s peak velocity sinusoidal, whole-body passive yaw rotations. Solid and dashed lines show first-order high-pass filter model fits to control and *Cyp26b1* cKO mouse population data, respectively. The high variability of phase and 0.02 Hz and relatively poor fit to gain data at 0.02Hz are due to the small amplitude of responses at that frequency. Differences between control and *Cyp26b1* cKO mice were not significant for G_A_, latency, model gain or model time constant (*P* < 0.05 for all comparisons). (**f**) Schematic view of apparatus for off-vertical axis rotation (OVAR). Pitch of the table was maintained at 17°. (**g**) No significant difference in the eye velocity was observed between control and *Cyp26b1* cKO mice during transient response (∼20 sec; amplitude: 43.8 ± 8.3 vs. 38.7 ± 7.4 deg (P = 0.66), time constant: 4.1 ± 1.1 vs. 5.1 ± 2.3 s (P = 0.68), when both semicircular canal and otolith organs are stimulated, and steady-state response (20 sec∼; amplitude: 5.3 ± 0.72 vs. 5.8 ± 0.75 deg (P = 0.64), bias: 10.5 ± 1.3 vs. 8.5 ± 1.3 (P = 0.33)), time when only responses from otolith organs are expected to be measured.

Next, to detect reflex responses derived from macular organs, off-vertical axis rotation (OVAR) testing was conducted^40^. For this purpose, we analyzed the recorded eye movements from the mice with heads fixed to a rotating platform (50 deg/s constant velocity), tilted 17 degrees with respect to the ground (Fig 6f). The eye velocity comprises two different responses: (1) an initial transient increase followed by a gradual exponential decrease and (2) steady-state response, in which eye velocity was oscillating around a constant bias with a sinusoidal waveform. We quantified the difference between the two groups by comparing the amplitude and time constant of the transient response as well as the amplitude and bias of the steady-state response. Similar to aVOR results, both transient steady-state OVAR responses were comparable for *Cyp26b1* cKO vs. controls (Fig. 6f, g). Together, these results indicate that despite the loss of striolar/central zone identity in vestibular sensory organs, horizontal aVOR and OVAR responses were normal in the *Cyp26b1* cKO mice, at least up to the maximal testable frequencies, which largely span the physiologically relevant range of head motion for mice in their natural environments^41^.

### *Cyp26b1* cKO mice performed better on rotarod but worse on balance beam tests

The vestibular system plays a vital role in balance control. To ensure stable body posture, vestibulo-spinal reflexes (VSR) produce compensatory movements of the neck and body to maintain the head in an upright position. Thus, we next tested *Cyp26b1* cKO mutants on a number of vestibular tasks used to test for balance and postural defects in mice. First, we found that these mice did not exhibit either circling or head tilting behaviors that are often associated with loss of vestibular function in rodents. Additionally, open-field testing revealed no hyperactivity in *Cyp26b1* cKO mutants (Supplemental video 1, Supplemental Fig.7), and the swimming ability of mutants was not affected under either light or dark conditions (data not shown). We further subjected animals to rotarod testing^42^ and quantified their performance based on the time the animal remained on the rotating rod with increasing acceleration. *Cyp26b1* cKO and *Foxg1^Cre^;Cyp26b1^lox/+^* controls exhibited similar motor performance on day 1 of the trial (Fig. 7a). However, *Cyp26b1* cKO mutants performed better (took longer time to fall) than controls on the second and third days (Fig. 7a). Whether this improved performance on the rotarod is due to replacement of striolar/central zone with extra-striolar/peripheral zone tissue or other mechanisms is not clear. Nevertheless, these collective behavioral results support the conclusion that the *Cyp26b1* cKO mutants exhibit no balance deficits during many of the tests commonly used to access balance and motor control functions in mice.

**Fig. 7.**
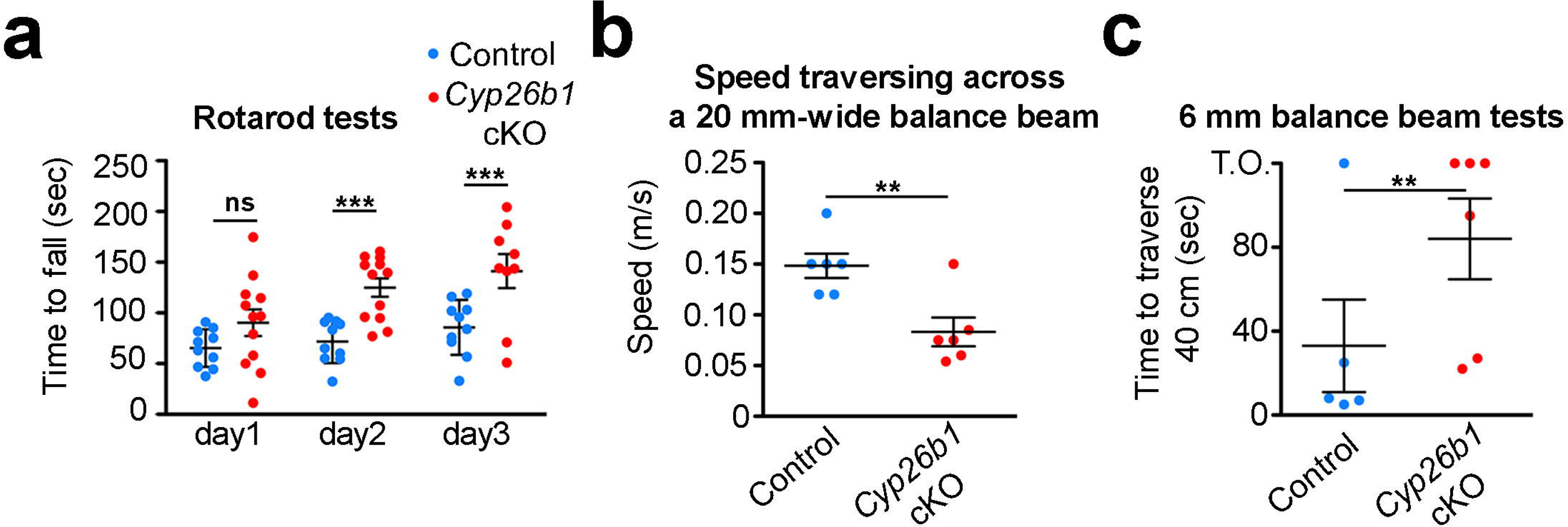
Impaired balance of *Cyp26b1* cKO mice during challenging vestibular-motor activities. (**a**) Quantification of rotarod tests. Each mouse was placed on a rotating rod, which accelerated from 5 to 40 rpm over a 5 min period. *Cyp26b1* cKO and *Foxg1^Cre^;Cyp26b1^lox/+^* controls exhibited similar motor performance on day 1 of the trial (90.3 ± 15.9 s in mutants, n = 10, vs. 65.3 ± 6.5 s in controls, n = 10, P = 0.2573). *Cyp26b1* cKO mutants were able to stay on rod longer than controls on the second day of testing (123.3 ± 11.0 s mutants vs. 71.7 ± 7.5 s controls, P = 0.0003) and third day (141.2 ± 17.8 s mutants vs 85.7 ± 9.5 s controls, P = 0.0037, Fig. 7a). (**b**) Quantification of speed to traverse 60 cm distance on a 20 mm-wide beam. *Cyp26b1* cKO mutants moved slower (0.083 ± 0.014 m/s, n = 6) than controls (0.148 ± 0.011 m/s, n = 5, P = 0.0056). (**c**) Quantification of time taken to cross 40 cm distance on a 6 mm-wide beam. Half of *Cyp26b1* cKO mutants failed to traverse the beam in 2 min (3/6), whereas most of controls reached the endpoints within 30 sec (4/5). By assigning 2 min for all the mice that failed to complete the 40 cm distance, *Cyp26b1* cKO mutants moved slower (taking 84.0 ± 19.24 s to traverse the beam) than controls (33.0 ± 22.04 s for controls, P = 0.0099). \*\**P* < 0.01 and \*\*\**P* < 0.001. T.O., time-out.

The balance beam test can detect subtle deficits in motor skills and balance that may not be detected by other standard motor and balance tests such as the rotarod test^43^. Accordingly, we tested the performance of *Cyp26b1* cKO mice on the balance beam. We found that *Cyp26b1* cKO mice walked slower than controls on a 20 mm wide beam over 60 cm length (Fig.7b). On a narrower beam (6 mm) over a distance of 40 cm, *Cyp26b1* cKO mice tended to freeze and stop walking (3 out of 6 vs. 1 out of 5 in controls) or moved slower than controls (Fig. 7c; Supplemental video 2). These results indicate that *Cyp26b1* cKO mice show a deficit in coordinating challenging vestibulomotor functions.

### Increased head tremor in *Cyp26b1* cKO mice

Despite normal aVOR, OVAR, and performance for all standard motor and balance tests other than balance beam testing, we observed that it was possible to distinguish *Cyp26b1* cKO mice from controls in their cages, because the former demonstrated head tremors. This distinctive feature of *Cyp26b1* cKO mice was particularly obvious at early postnatal ages (Supplemental video 3). Quantitative analysis at P9 showed that the incidence of head tremors per 100 mm distance travelled was higher in mutants than controls (Fig. 8a) and each head tremor episode also lasted longer in mutants (Fig. 8b). These head tremors became less noticeable during normal activities in a cage as the mutants matured (Supplemental video 1), suggesting some level of compensation. To quantify head tremors in adult *Cyp26b1* cKO mutants, we affixed a 6D MEMS module consisting of three gyroscopes and three linear accelerometers (see Materials and Methods) to the mouse’s head implant. We confirmed that, when mice were not actively moving through their environment, *Cyp26b1* cKO adults exhibit head tremors (Fig. 8c, d). Power spectra of head movements revealed higher power for *Cyp26b1* cKO mutants than controls at high frequency (5-20 Hz) for all six dimensions (Fig. 8e, f). For mid-frequency (1-5 Hz), *Cyp26b1* cKO mutants have higher power for yaw and roll head velocity (Fig. 8f). Taken together, these results indicate that the ability to maintain head stabilization in *Cyp26b1* cKO mice is compromised.

**Fig. 8.**
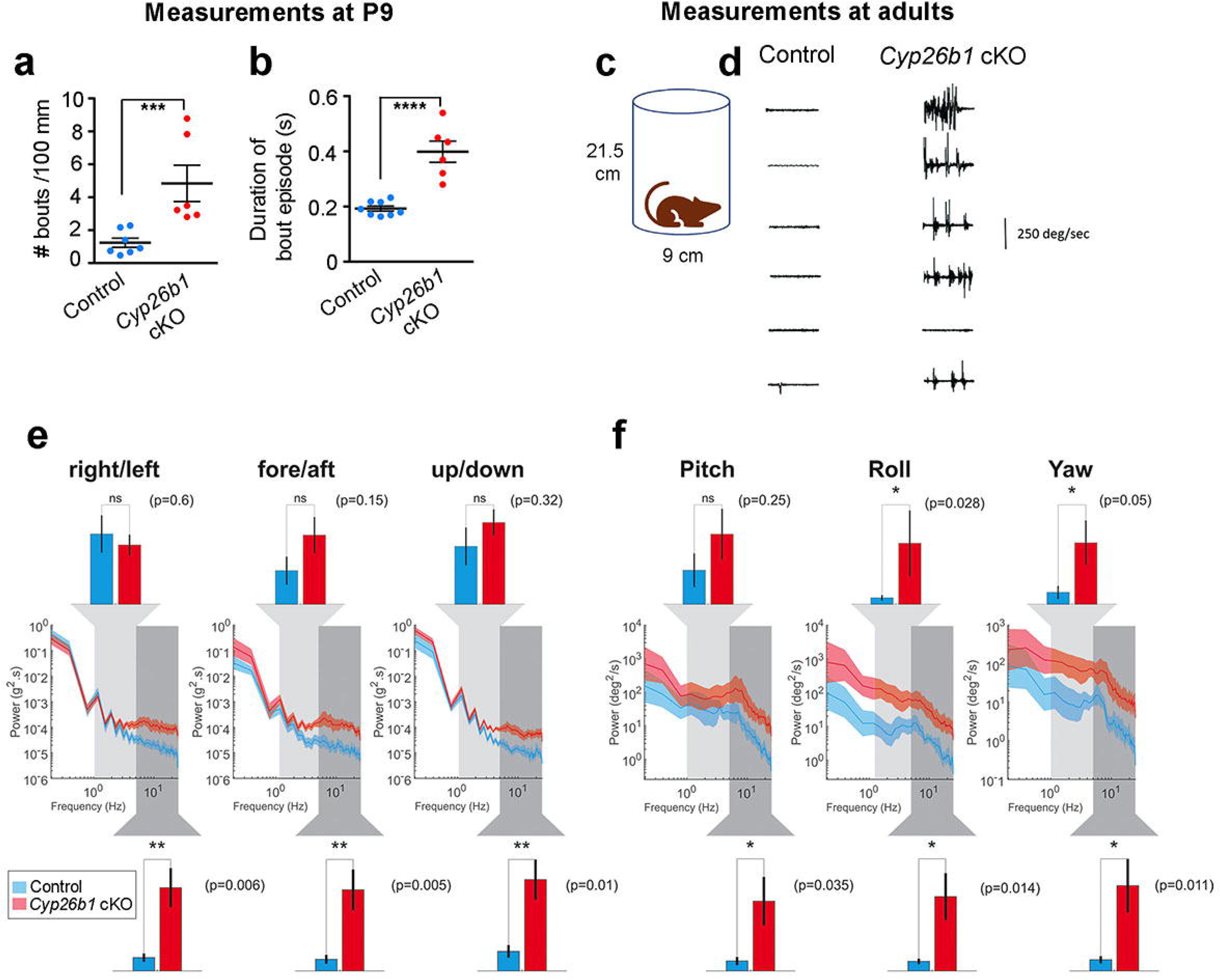
Increased head tremors in *Cyp26b1* cKO mice. (**a**, **b**) Quantification of head-tremors for P9 pups. The number of bouts normalized over a 100 mm distance traveled (**a**) and average duration of each head-tremor episode over a 10-min period are shown (**b**). *Cyp26b1* cKO pups showed more frequent (**a**, 4.8 ± 1.1 in mutants, n = 6, vs. 1.2 ± 0.2 in controls, n = 8, P = 0.0059) and longer duration (**b**, 398 ± 38 ms in mutants vs. 192 ± 9 ms in controls, n = 8, P < 0.0001) head tremors, compared to *Foxg1^Cre^;Cyp26b1^lox/+^* controls. (**c**) Schematic of the apparatus for measuring head tremors of an adult mouse at rest with a miniture head motion sensor affixed on the top of the skull. (**d**) *Cyp26b1* cKO showed characteristic head tremors (5/6), which were not present in controls. (**e**, **f**) Comparison of power spectra density of head movements in translational axes (**e**) and rotational axes (**f**) between controls (blue) and *Cyp26b1* cKO mutants (red). *Cyp26b1* cKO exhibit siginificantly higher power than controls at high frequencies (5-20 Hz, P = 0.006 for right/left, P = 0.005 for fore/aft, P = 0.01 for up/down, P = 0.035 for pitch, P = 0.014 for roll, and P = 0.011 for yaw axis, n = 6/group). Angular head velocity for yaw and roll axes of *Cyp26b1* cKO also had significantly higher power for frequencies between 1 and 5 Hz (upper graphs, P = 0.05 for yaw, and P = 0.028 for roll axis). Unpaired t-test was applied for all tests. \**P* < 0.05. \*\**P* < 0.01 and \*\*\**P* < 0.001.

## DISCUSSION

### Striolar/central zones of vestibular organs require reduced RA signaling

Based on molecular, cellular and physiological evidence, we demonstrated that striolar/central zones of vestibular sensory organs in the *Cyp26b1* mutants (gain-of RA signaling) are severely reduced. We concluded that reduced RA signaling mediated by the expression of *Cyp26b1* in the prospective striolar/central zones is required for this regional formation. This conclusion is supported by results from a complementary mouse model, *Aldh1a3^-/-^* mice, in which endogenous RA signaling is reduced in the peripheral region of sensory organs and striolar HCs are expanded in the two maculae (Fig. 2, Supplemental Fig. 2). The expansion of Ocm*^+^* HC in the central zone is not observed in the cristae (Supplemental Fig. 3). We attributed this difference between maculae and cristae to possible functional redundancy provided by *Aldh1a1* and *Aldh1a2,* which are expressed in the adjacent nonsensory tissues or in the roof of the sensory organs^44^. Additionally, the extent of expansion of Ocm^+^ HCs is different between the utricle and saccule (Fig. 2c and S2c). Therefore, it is possible that intrinsic molecular differences across different regions of a sensory organ could also affect the response to the loss of RA signaling.

It is unclear how lower RA signaling mediates the unique cellular and anatomical features of the central regions during development. Given that expression of both *Cyp26b1* and *Aldh1a3* is concentrated in the SCs (Fig. 1f, g), it is possible that RA levels regulated by SCs provide a niche for the HCs and afferent nerve endings contained within the sensory epithelium. Receptors of RA are known to be expressed in the otic vesicle and sensory epithelium of the inner ear^45–47^. Alternatively, the reduced RA in striolar SCs could regulate downstream genes within SC, which indirectly affect the development of HCs and their afferents. Previous results have suggested such a function for SCs^48^. Activated erbB receptors in SCs of the utricle lead to production of BDNF, which promotes synaptogenesis between HCs and afferent neurons.

### The striola generates the vestibular evoked potential (VsEP)

VsEP measure utricular and saccular function and are produced in response to transient changes in the linear head acceleration (jerk)^17, 18^. Several lines of evidence have implicated the irregular afferents of the striola in the generation of VsEP. First, the phase-locking and transient firing properties of irregular neurons render them the most likely candidates to respond in a synchronized manner to linear jerk stimuli and produce the signature compound action potentials^2, 20^. Second, VsEP is impaired by inhibitors of KCNQ, which are subunits of potassium channels that are most abundant in the calyceal endings of the striola^19^. Third, the linear pulse stimuli used to evoke VsEP are likely to activate irregular, striolar, afferents based on experiments with sound and bone vibration stimuli in the 500-1000 Hz range^20, 49^.

Our results provide strong evidence that the VsEP originates in the striola. The loss of morphological and physiological features in the striola of *Cyp26b1* cKO mice together with the VsEP suggests that the striola is necessary for the generation of VsEP. Earlier it was established that otoconia are essential for generating VsEP^17, 50^; here we show that the presence of otoconia is not sufficient for VsEP. In *Cyp26b1* cKO mice, otoconia are present but have extrastriolar-like properties throughout the epithelium. In control otolith organs, mechanical input to HCs in different zones is differentially shaped by striking differences in otoconia, otolithic gel layers, stereocilia- bundle morphology and bundle coupling to the otoconia^7^. These differences are likely to contribute to the greater sensitivity of striolar afferents to high-frequency and transient head motion, including the linear pulse stimuli used to evoke VsEPs. Other likely factors are the zonal differences in calcium binding proteins, afferent calyceal synapses and excitability, all absent in *Cyp26b1* cKO.

### Striolar/central zones are not essential for vestibulo-ocular reflexes

Given the larger cell bodies, higher conduction velocities^51, 52^ and better phase-lead properties^53–55^ of the irregular neurons in the striolar/central zones than regular neurons in the extra-striola/peripheral zones, these regions have been postulated to be important for mediating fast vestibular reflexes^56^. The aVOR in mice is fast with a latency as low as < 7 milliseconds^57^. However, we did not detect any deficit in aVOR and OVAR in *Cyp26b1* cKO mutants when lateral cristae and macular organs were inertially stimulated respectively, suggesting that the striolar/central zone of vestibular organs (preferentially innervated by irregular afferents) are dispensable for these functions, at least in the adults. We cannot rule out the possibility that vestibular reflexes at a younger age or stimulations beyond frequencies tested (10 Hz in the case of aVOR) may require normal function of the central zones. These results further suggest that regular afferents concentrated in the extrastriolar/peripheral zones of the vestibular organs are important for mediating VOR. It was reported that *Aldh1a3^-/-^* (em RA) mice show absence of aVOR and OVAR responses^31^, and our results show that the striolar/central zones in the vestibular organs are expanded based on Ocm staining (Fig. 2). Thus, these results are consistent with the notion that the extrastriolar/peripheral zones are more important than striolar/central zones for aVOR and OVAR. When irregular afferents in squirrel monkeys were selectively affected using galvanic currents, aVOR was not changed^58^. Together, these results suggest that regular afferents in the extrastriolar/peripheral zone rather than irregular afferents in the striolar/central zone are more important for mediating angular and linear VOR in mice.

### *Cyp26b1* cKO mice show deficits in precise balance control and self-motion

A long-standing view is that irregular afferents, which innervate the striolar/central zone, preferentially contribute to VSR^59, 60^. Our finding that loss of striolar/central zones leads to a deficit in coordinating challenging postural control on the balance beam is consistent with this proposal^59, 60^. Surprisingly, however, *Cyp26b1* cKO mice performed well on other less sensitive tests commonly used for accessing changes in balance and motor control function in vestibularly deficient mice. Further, we did not observe classical vestibular behavioral deficits such as head tilt and circling in the *Cyp26b1* cKO mutants. Together these two observations suggest that the extrastriolar/peripheral zones, which constitute 75-80% of the sensory epithelia^4, 5, 11, 12^ and are innervated by regular afferents, play a critical role in vestibulo-spinal as well as vestibulo-ocular reflexes in mice. In support of this hypothesis, regular afferents with their sustained firing properties (less adapting) are thought to be more important in conveying head positional information in steady state such as head tilts^61^.

A distinguishing characteristic of *Cyp26b1* cKO mice was that they demonstrated head tremor. Similar head movement behaviors have been reported in both primates and humans with compromised vestibular functions. For example, squirrel monkeys show transient head tremor and postural instability after plugging one of the lateral semicircular canals^62^. Patients with chronic bilateral vestibular loss that exhibit gaze variability and oscillopsia also show pronounced head oscillations in response to weighted head mass, compared to healthy individuals^63^. This head tremor behavior is attributed to a failure of vestibular input that is normally required to maintain head stability^64, 65^ and ensure movement accuracy during active goal-directed behaviors^63, 66^. It is tempting to speculate that in the *Cyp26b1* cKO mouse, poor or abnormal vestibular inputs from the striolar/central regions cause head tremors or oscillations, which affected challenging vestibulo-motor activities such as traversing a narrow balance beam when rapid vestibular input to motor centers is required. One caveat is that *Cyp26b1* cKO mice show some loss of *Cyp26b1* expression in the brain (data not shown), suggesting that the behavioral phenotype may result in part from central as well as peripheral *Cyp26b1* loss.

In summary, using a genetic approach we generated a viable mouse mutant that largely lacks the striolar/central zone of vestibular organs. By disrupting RA signaling during embryogenesis, the entire axis of this specialized zone failed to develop properly, including associated components such as the otoconia and innervating neurons. As a result, the loss of this highly specialized and conserved region^67^ selectively affected the ability of afferents to respond to transient changes in linear acceleration and may affect vestibular input. Interestingly, low levels of RA signaling mediated by two of the RA degradation enzymes, Cyp26a1 and Cyp26c1, are also required for the formation of the fovea, a high acuity area of the retina^23^. Therefore, the same developmental strategy is being employed to generate complexity and specialization in two different sensory systems. A better understanding of the striolar/central zone-specific function has clinical and therapeutic relevance as HCs in this region are more susceptible to ototoxic insults in animal models^68^.

## METHODS

All of the methods are described in the supplementary information.

## ACKNOWLEDGEMENTS

We would like to thank Drs. Yogita Chudasama and Johann du Hoffmann and Mr. Kevin Cravedi in the Mouse Behavioral Core at NIMH, Drs. Tracy Fitzgerald and Talah Wafa in the Mouse Auditory Testing Core Facility at NIDCD for their advice and technical assistance in behavioral and VsEP testing. We also thank Dr. Ronald S. Petralia in Advanced Imaging Core at NIDCD for technical assistance on SEM. We are also grateful to Drs. Lisa Cunningham and Thomas Friedman and members of the Wu Lab for their critical review of the manuscript. This work was funded by Intramural Research Program grant (#1ZIADC000021) to DKW, NIH R01 grants DCO2290 and DC012347 to RAE, NIH RO1 grants DC002390 and R01-DC013069 and Canadian Institutes of Health Research grant to KEC.

## REFERENCES

1. Ahmed, R. M., Hannigan, I. P., MacDougall, H. G., Chan, R. C. & Halmagyi, G. M. Gentamicin ototoxicity: a 23-year selected case series of 103 patients. The Medical Journal of Australia 196, 701–704, doi:10.5694/mja11.10850 (2012).

2. Jones, T. A., Lee, C., Gaines, G. C. & Grant, J. W. On the high frequency transfer of mechanical stimuli from the surface of the head to the macular neuroepithelium of the mouse. J Assoc Res Otolaryngol 16, 189–204, doi:10.1007/s10162-014-0501-9 (2015).

3. Eatock, R. A. & Songer, J. E. Vestibular hair cells and afferents: two channels for head motion signals. Annu Rev Neurosci 34, 501–534, doi:10.1146/annurev-neuro-061010-113710 (2011).

4. Desai, S. S., Zeh, C. & Lysakowski, A. Comparative morphology of rodent vestibular periphery. I. Saccular and utricular maculae. J Neurophysiol 93, 251–266, doi:10.1152/jn.00746.2003 (2005).

5. Desai, S. S., Ali, H. & Lysakowski, A. Comparative morphology of rodent vestibular periphery. II. Cristae ampullares. J Neurophysiol 93, 267–280, doi:10.1152/jn.00747.2003 (2005).

6. Lysakowski, A. et al. Molecular microdomains in a sensory terminal, the vestibular calyx ending. J Neurosci 31, 10101–10114, doi:10.1523/JNEUROSCI.0521-11.2011 (2011).

7. Lim, D. J. Otoconia in health and disease. A review. Ann. Otol. Rhinol. Laryngol. Suppl., 17–24 (1984).

8. Li, A., Xue, J. & Peterson, E. H. Architecture of the mouse utricle: macular organization and hair bundle heights. J Neurophysiol 99, 718–733, doi:10.1152/jn.00831.2007 (2008).

9. Fernandez, C., Goldberg, J. M. & Baird, R. A. The Vestibular Nerve of the Chinchilla III. Peripheral Innervation Patterns in the Utricular Macula. Journal of Neurophysiology 63, 767–780 (1990).

10. Fernandez, C., Baird, R. A. & Goldberg, J. M. The Vestibular Nerve of the Chinchilla. I. Peripheral Innervation Patterns in the Horizontal and Superior Semicircular Canals. Journal of Neurophysiology 60, 167–181 (1988).

11. Goldberg, J. M., Desmadryl, G., Baird, R. A. & Fernandez, C. The Vestibular Nerve of the Chinchilla. V. Relation Between Afferent Discharge Properties and Peripheral Innervation Patterns in the Utricular Macula. Journal of Neurophysiology (1990).

12. Baird, R. A., Desmadryl, G., Fernandez, C. & Goldberg, J. M. The Vestibular Nerve of the Chinchilla. II. Relation Between Afferent Response Properties and Peripheral Innervation Patterns in the Semicircular Canals. Journal of Neurophysiology 60, 182–203 (1988).

13. Eatock, R. A. Specializations for Fast Signaling in the Amniote Vestibular Inner Ear. Integr Comp Biol 58, 341–350, doi:10.1093/icb/icy069 (2018).

14. Kalluri, R., Xue, J. & Eatock, R. A. Ion channels set spike timing regularity of mammalian vestibular afferent neurons. J Neurophysiol 104, 2034–2051, doi:10.1152/jn.00396.2010 (2010).

15. Jamali, M., Chacron, M. J. & Cullen, K. E. Self-motion evokes precise spike timing in the primate vestibular system. Nat Commun 7, 13229, doi:10.1038/ncomms13229 (2016).

16. Jones, T. A. et al. The adequate stimulus for mammalian linear vestibular evoked potentials (VsEPs). Hear Res 280, 133–140, doi:10.1016/j.heares.2011.05.005 (2011).

17. Jones, S. M., Erway, L. C., Johnson, K. R., Yu, H. & Jones, T. A. Gravity receptor function in mice with graded otoconial deficiencies. Hear Res 191, 34–40, doi:10.1016/j.heares.2004.01.008 (2004).

18. Dulon, D., Safieddine, S., Jones, S. M. & Petit, C. Otoferlin is critical for a highly sensitive and linear calcium-dependent exocytosis at vestibular hair cell ribbon synapses. J Neurosci 29, 10474–10487, doi:10.1523/JNEUROSCI.1009-09.2009 (2009).

19. Lee, C., Holt, J. C. & Jones, T. A. Effect of M-current modulation on mammalian vestibular responses to transient head motion. J Neurophysiol 118, 2991–3006, doi:10.1152/jn.00384.2017 (2017).

20. Curthoys, I. S., MacDougall, H. G., Vidal, P. P. & de Waele, C. Sustained and Transient Vestibular Systems: A Physiological Basis for Interpreting Vestibular Function. Front Neurol 8, 117, doi:10.3389/fneur.2017.00117 (2017).

21. Rhinn, M. & Dolle, P. Retinoic acid signalling during development. Development 139, 843–858, doi:10.1242/dev.065938 (2012).

22. Bok, J. et al. Transient retinoic acid signaling confers anterior-posterior polarity to the inner ear. PNAS 108, 161–166 (2011).

23. da Silva, S. & Cepko, C. L. Fgf8 Expression and Degradation of Retinoic Acid Are Required for Patterning a High-Acuity Area in the Retina. Dev Cell 42, 68–81 e66, doi:10.1016/j.devcel.2017.05.024 (2017).

24. Dubey, A., Rose, R. E., Jones, D. R. & Saint-Jeannet, J. P. Generating retinoic acid gradients by local degradation during craniofacial development: One cell’s cue is another cell’s poison. Genesis 56, doi:10.1002/dvg.23091 (2018).

25. Cunningham, T. J. & Duester, G. Mechanisms of retinoic acid signalling and its roles in organ and limb development. Nat Rev Mol Cell Biol 16, 110–123, doi:10.1038/nrm3932 (2015).

26. Rau, A., Legan, P. K. & Richardson, G. P. Tectorin mRNA Expression Is Spatially and Temporally Restricted During Mouse Inner Ear Development. The Journal of Comparative Neurology 405, 271–280 (1999).

27. Simmons, D. D., Tong, B., Schrader, A. D. & Hornak, A. J. Oncomodulin identifies different hair cell types in the mammalian inner ear. J Comp Neurol 518, 3785–3802, doi:10.1002/cne.22424 (2010).

28. Yashiro, K., et al. Regulation of Retinoic Acid Distribution Is Required for Proximodistal Patterning and Outgrowth of the Developing Mouse Limb. (2004).

29. Dupe, V. et al. A newborn lethal defect due to inactivation of retinaldegyde dehydrogenase type 3 is prevented by maternal retinoic acid treatment. PNAS 100, 14036–14041 (2003).

30. Hebert, J. M. & McConnell, S. K. Targeting of cre to the Foxg1 (BF-1) locus mediates loxP recombination in the telencephalon and other developing head structures. Dev Biol 222, 296–306, doi:10.1006/dbio.2000.9732 (2000).

31. Romand, R. et al. Retinoic acid deficiency impairs the vestibular function. J Neurosci 33, 5856–5866, doi:10.1523/JNEUROSCI.4618-12.2013 (2013).

32. Leonard, R. B. & Kevetter, G. A. Molecular probes of the vestibular nerve I. Peripheral termination patterns of calretinin. Brain Research 928, 8–17 (2002).

33. Deans, M. R. et al. Asymmetric distribution of prickle-like 2 reveals an early underlying polarization of vestibular sensory epithelia in the inner ear. J Neurosci 27, 3139–3147, doi:10.1523/JNEUROSCI.5151-06.2007 (2007).

34. Jiang, T., Kindt, K. & Wu, D. K. Transcription factor Emx2 controls stereociliary bundle orientation of sensory hair cells. Elife 6, doi:10.7554/eLife.23661 (2017).

35. Meredith, F. L. & Rennie, K. J. Channeling your inner ear potassium: K(+) channels in vestibular hair cells. Hear Res 338, 40–51, doi:10.1016/j.heares.2016.01.015 (2016).

36. Contini, D., Price, S. D. & Art, J. J. Accumulation of K(+) in the synaptic cleft modulates activity by influencing both vestibular hair cell and calyx afferent in the turtle. J Physiol 595, 777–803, doi:10.1113/JP273060 (2017).

37. Songer, J. E. & Eatock, R. A. Tuning and timing in mammalian type I hair cells and calyceal synapses. J Neurosci 33, 3706–3724, doi:10.1523/JNEUROSCI.4067-12.2013 (2013).

38. Iwasaki, S., Chihara, Y., Komuta, Y., Ito, K. & Sahara, Y. Low-voltage-activated potassium channels underlie the regulation of intrinsic firing properties of rat vestibular ganglion cells. J Neurophysiol 100, 2192–2204, doi:10.1152/jn.01240.2007 (2008).

39. Stahl, J. S. Using eye movements to assess brain function in mice. Vision Res 44, 3401–3410, doi:10.1016/j.visres.2004.09.011 (2004).

40. Beraneck, M., Bojados, M., Le Seac’h, A., Jamon, M. & Vidal, P. P. Ontogeny of mouse vestibulo-ocular reflex following genetic or environmental alteration of gravity sensing. PLoS One 7, e40414, doi:10.1371/journal.pone.0040414 (2012).

41. Carriot, J., Jamali, M., Chacron, M. J. & Cullen, K. E. The statistics of the vestibular input experienced during natural self-motion differ between rodents and primates. J Physiol 595, 2751–2766, doi:10.1113/JP273734 (2017).

42. Tung, V. W., Burton, T. J., Dababneh, E., Quail, S. L. & Camp, A. J. Behavioral assessment of the aging mouse vestibular system. J Vis Exp, doi:10.3791/51605 (2014).

43. Luong, T. N., Carlisle, H. J., Southwell, A. & Patterson, P. H. Assessment of motor balance and coordination in mice using the balance beam. J Vis Exp, doi:10.3791/2376 (2011).

44. Romand, R. et al. Dynamic expression of retinoic acid-synthesizing and - metabolizing enzymes in the developing mouse inner ear. J Comp Neurol 496, 643–654, doi:10.1002/cne.20936 (2006).

45. Dolle, P., Fraulob, V., Kastner, P. & Chambon, P. Developmental expression of murine retinoid X receptor (RXR) genes. Mechanisms of Development 45, 91–104 (1994).

46. Romand, R. et al. The retinoic acid receptors RARa and RARg are required for inner ear development. Mechanisms of Development 119, 213–223 (2002).

47. Shen, J., Scheffer, D. I., Kwan, K. Y. & Corey, D. P. SHIELD: an integrative gene expression database for inner ear research. Database (Oxford) 2015, bav071, doi:10.1093/database/bav071 (2015).

48. Gomez-Casati, M. E. et al. Nonneuronal cells regulate synapse formation in the vestibular sensory epithelium via erbB-dependent BDNF expression. PNAS 107, 17005–17010 (2010).

49. McCue, M. P. & Guinand, J. J. Acoustically Responsive Fibers in the Vestibular Nerve of the Cat. Journal of Neuroscience 14, 6058–6070 (1994).

50. Jones, T. A., Jones, S. M. & Hoffman, L. F. Resting discharge patterns of macular primary afferents in otoconia-deficient mice. J Assoc Res Otolaryngol 9, 490–505, doi:10.1007/s10162-008-0132-0 (2008).

51. Lysakowski, A., Minor, L. B., Fernandez, C. & Goldberg, J. M. Physiological Identification of Morphologically Distinct Afferent Classes Innervating the Cristae Ampullares of the Squirrel Monkey. Journal of Neurophysiology 73, 1270–1281 (1995).

52. Goldberg, J. M. & Fernandez, C. Conduction times and background discharge of vestibular afferents. Brain Research 122, 545–550 (1977).

53. Hullar, T. E. et al. Responses of irregularly discharging chinchilla semicircular canal vestibular-nerve afferents during high-frequency head rotations. J Neurophysiol 93, 2777–2786, doi:10.1152/jn.01002.2004 (2005).

54. Kim, K. S., Minor, L. B., Della Santina, C. C. & Lasker, D. M. Variation in response dynamics of regular and irregular vestibular-nerve afferents during sinusoidal head rotations and currents in the chinchilla. Exp Brain Res 210, 643–649, doi:10.1007/s00221-011-2600-8 (2011).

55. Sadeghi, S. G., Minor, L. B. & Cullen, K. E. Response of vestibular-nerve afferents to active and passive rotations under normal conditions and after unilateral labyrinthectomy. J Neurophysiol 97, 1503–1514, doi:10.1152/jn.00829.2006 (2007).

56. Huterer M. Cullen, K. E. Vestibuloocular Reflex Dynamics During High-Frequency and High-Acceleration Rotations of the Head on Body in Rhesus Monkey. J Neurophysiol 88, 13–28 (2002).

57. Hubner, P. P., Khan, S. I., Lasker, D. M. & Migliaccio, A. A. Core Body Temperature Effects on the Mouse Vestibulo-ocular Reflex. J Assoc Res Otolaryngol 18, 827–835, doi:10.1007/s10162-017-0639-3 (2017).

58. Minor, L. B. & Goldberg, J. M. Vestibular-Nerve Inputs to the Vestibulo-ocular Reflex: A Functional-Ablation Study in the Squirrel Monkey. J Neurosci 11, 1636–1648 (1991).

59. Goldburg, J. M. & Fernandez, C. Physiology of Peripheral Neurons Innervating Semicircular Canals of the Squirrel Monkey. III. Variations Among Units in Their Discharge Properties. J Neurophysiol 34, 676–684 (1971).

60. Bilotto, G., Goldberg, J. M., Peterson, B. W. & Wilson, V. J. Dynamic Properties of Vestibular Reflexes in the Decerebrate Cat. Exp Brain Res 47, 343–352 (1982).

61. Fernandez, C. & Goldberg, J. M. Physiology of Peripheral Neurons Innervating Otolith Organs of the Squirrel Monkey. I. Response to Static Tilts and to Long-Duration Centrifugal Force. Journal of Neurophysiology 39, 970–984 (1976).

62. Paige, G. D. Vestibuloocular Reflex and Its Interactions With Visual Following Mechanisms in the Squirrel Monkey. II. Response Characteristics and Plasticity Following Unilateral Inactivation of Horizontal Calanl. Journal of Neurophysiology 49, 152–168 (1983).

63. Saglam, M., Glasauer, S. & Lehnen, N. Vestibular and cerebellar contribution to gaze optimality. Brain 137, 1080–1094, doi:10.1093/brain/awu006 (2014).

64. Goldberg, J. M. & Cullen, K. E. Vestibular control of the head: possible functions of the vestibulocollic reflex. Exp Brain Res 210, 331–345, doi:10.1007/s00221-011-2611-5 (2011).

65. Angelaki, D. E. & Cullen, K. E. Vestibular system: the many facets of a multimodal sense. Annu Rev Neurosci 31, 125–150, doi:10.1146/annurev.neuro.31.060407.125555 (2008).

66. Sylvestre, P. A. & Cullen, K. E. Premotor correlates of integrated feedback control for eye-head gaze shifts. J Neurosci 26, 4922–4929, doi:10.1523/JNEUROSCI.4099-05.2006 (2006).

67. Lysakowski, A. & Goldberg, J. M. Morphophysiology of the Vestibular Periphery. The Vestibular System, 57–152 (2004).

68. Lindeman, H. H. Regional Differences in Sensitivity of the Vestibular Sensory Epithelia to Ototoxic Antibiotics. Acta Oto-Laryngologica 67, 177–189, doi:10.3109/00016486909125441 (2009).

69. Spoon, C., Moravec, W. J., Rowe, M. H., Grant, J. W. & Peterson, E. H. Steady-state stiffness of utricular hair cells depends on macular location and hair bundle structure. J Neurophysiol 106, 2950–2963, doi:10.1152/jn.00469.2011 (2011).

