## Supplemental file for "Cytochrome P450 26b1-mediated specification of vestibular striola and central zones is required for transient responses in linear acceleration"

**Supplementary information**

**MATERIALS & METHODS**

**Mice.** The following mouse strains were used in the study: *Aldh1a3^+/-^* (maintained in a mixed C57BL/6J and CD1 background)^1^, *Cyp26b^flox/flox^* (RIKEN BRC (RBRC04333), maintained in a C57BL/6J background) and *Foxg1^Cre^*^2^ (RRID:[IMSR_JAX:004337](https://scicrunch.org/resolver/IMSR_JAX:004337), maintained in C57BL/6J background). In addition, *Cyp26b1^+/-^* mice were generated by breeding *Cyp26b1^flox/flox^* mice with a ubiquitous cre line, *Actin^Cre^*, and maintained in a C57BL/6J background after recombination and removal of the *cre* allele. *Foxg1^Cre^; Cyp26b1^flox/-^* were produced by crossing *Foxg1^Cre^;Cyp26b1^+/-^* males with *Cyp26b1^flox/flox^* females. All animal experiments were conducted under the approved NIH animal protocols at the NIH, University of Chicago, University of Nebraska -Lincoln, Johns Hopkins University and according to NIH animal user guidelines.

**Tissue preparation.** Timed pregnant females or postnatal mice were harvested and whole heads were hemi-sected, brain is removed and fixed in 4% paraformaldehyde overnight. Then, half-heads were cryo-preserved and stored in -80^o^C until subsequently processed for cryo-sectioning or whole mount dissection.

***In situ* hybridization.** *In situ* hybridization was conducted as previously described^3^. Digoxigenin-labeled RNA probes were generated for *Cyp26b1* (GenBank: AW049789), *β-tectorin*^4^, *Aldh1a3^5^* as described.

**Whole-mount immunohistochemistry.** Dissected saccules or utricles with anterior cristae and lateral cristae attached were blocked with PBS containing 4% normal donkey serum and 0.2% Triton X (PBT). Then, specimens were treated with primary antibodies diluted with blocking solution overnight at 4°C. The primary antibodies used were as follow: goat polyclonal anti-oncomodulin (1:300, Santa Cruz Biotech, #sc-7446), rabbit polyclonal anti-β-tectorin (1:1000, a gift from Guy Richardson, U of Sussex), rabbit polyclonal anti-Myosin7a (1:1000, Proteus, #25-6790), rabbit polyclonal anti-Calbindin (1:1000, Millipore, #AB1778), mouse monoclonal anti-βIII tubulin (1:500; R&D, #MAB1195), mouse anti-βII-spectrin (1:500; BD Bioscience, #612562), goat polyclonal anti-Sox2 (1:500; Santa Cruz Biotech, #sc-17320), and rabbit polyclonal anti-Arl13b (1:1000; Abcam, #83879). Alexa Fluor 647-conjugated phalloidin was used to label actin-based stereocilia (Thermo Fisher Scientific, #A22287).

Following primary antibody incubation, samples were washed with PBT extensively before incubating with appropriate secondary antibodies conjugated with fluorescent proteins: donkey anti-mouse, rabbit, or goat IgG (H+L) antibody (Thermo Fisher Scientific) for 1 hour at 4^o^C. Then, samples were washed extensively with PBT before mounting with ProLong Gold Antifade (Invitrogen) and imaged with a Zeiss LSM780 confocal microscope. All low-power immunostaining pictures are composite of images taken at 40x magnification.

**Scanning electron microscopy (SEM).** The preparation of samples for SEM was conducted as described^6^. Briefly, the utricles and cristae were quickly dissected from harvested animals and submerged in fresh fixative consisting of 2.5% glutaraldehyde (Electron Microscopy Sciences), 2% formaldehyde (EMS), 3 mM calcium chloride, and 0.1 M cacodylate. Two hours after fixation at room temperature, the otoconia was exposed and then tissues were post-fixed with osmium-thiocarbohydrazide-osmium (OTOTO) method. Specimens were then dehydrated with a series of ethanol followed by critical point drying. After spattering with platinum for coating, pictures were taken using electron microscope (SU4800, Hitachi).

**Measurement of otoconial size.** SEM images were used to measure the length of each otoconia in control and mutant utricles (n = 2 for each genotype). To identify the central region in *Cyp26b1* cKO, the striola was identified in control utricles based on the smaller size of the otoconia. Then, a comparable region in the two mutant utricles was compared to that of the controls. The length of the otoconial crystals in the middle of striolar (4 mm^2^) and medial extrastriolar region were measured by using ImageJ software.

**Measurement of K/S ratio.** P10 utricle samples (n = 4 for each genotype) were fixed and labeled with conjugated phalloidin, anti-Arl13b (for kinocilium) and anti-β-spectrin antibodies. A composite picture of confocal images taken at 40x magnification was generated and LPR was drawn based on hair bundle orientations. Then, regions lateral (lateral extrastriola) and immediately medial (striola) to the LPR in the center of the utricle were selected for confocal images taken at 63x magnification and used for the kinocilium and the tallest sterocilium measurements using Volocity (PerkinElmer) or imageJ software. Ratio of the height of the kinocilium (K) to the height of the tallest stereocilia (S) was calculated for 3-5 HCs in each region.

**Retinoic acid treatment of timed pregnant females.** Viable *Aldh1a3^-/-^* mice was generated as described^7^. Briefly, retinoic acid powder (Sigma) was suspended in ethanol (5 mg/ml), and then 1 ml of RA solution was mixed with 50 mg of normal chow (5015, LabDiet) and administered to pregnant females *ad libitum* from E8.5 to E14.5.

**Measurement of HC density.** HC density in the utricle was measured as described^8^. Briefly, a straight line across the widest region of an utricle was drawn along the anterior-posterior (A-P) axis. Then, two lines perpendicular to the A-P line, which mark the middle-third region of the utricle were drawn. This middle region was divided into two equal halves along the medial-lateral axis, and the posterior half was further sub-divided into 4 equal regions marked as 1, 2, 3, and 4, representing LES, striola, and two MES regions, respectively (Supplemental Fig. 4a). The number of HCs per 0.01 mm^2^ area within areas 1 and 2 were counted (n = 4).

**Quantification of complex calyces.** Wholemount utricles of *Cyp26b1* cKO and littermate controls at P30-45 were immunolabelled with anti-Tuj1 and anti-calbindin antibodies (n = 3/group). Z-stacked images were taken using a laser scanning confocal microscope (Zeiss LSM780). Tuj1-positive calyces that surround two or three HC bodies in each utricle were scored manually by examining individual confocal stacks.

**Whole-cell patch clamp recordings**. Data shown here are from 21 littermates: 11 control mice (*Foxg1^Cre^;Cyp26b1^lox/+^*, ages P17-P97, median P21) and 10 mutant mice (*Cyp26b1* cKO , n = 10, P12-P100, median P20.5). Tissue preparation was conducted as described^9^. For each electrophysiological experiment, a mouse was anesthetized deeply by exposure to isoflurane and decapitated. The utricle plus the superior division of the vestibular ganglion and distal part of the vestibular nerve were excised, trimmed, and secured in a recording chamber with the exposed apical surface of the epithelium facing up, as described^9,10^.

*Recordings.* Whole-cell recordings were made at room temperature (23-25^o^C) using pipettes with resistances between 3 and 7 M$\Omega$ in standard solutions. The bath (external) solution was Leibovitz-15 (L15) medium, supplemented with 10 mM HEPES (4-(2-Hydroxyethyl)piperazine-1-ethanesulfonic acid, N-(2-Hydroxyethyl)piperazine-N′-(2-ethanesulfonic acid), ~315 mmol kg^–1^ and pH 7.4. The pipette (internal) solution comprised (in mM): 135 KCl, 0.1 CaCl_2_, 3.5 MgCl_2_, 3 Na_2_ATP, 5 creatine phosphate (Na^+^ salt), 0.1 Na-cAMP, 0.1 Li-GTP, 5 EGTA, and 5 HEPES. The solution was brought to pH 7.3 and ~300 mmol kg^–1^ by adding ~28 mM KOH. Sulforhodamine 101 (1 mg/100 ml; Invitrogen) was added to the internal solution to label the recorded HC or calyx.

Whole-cell, GΩ-seal recordings were made from visually identified HCs or calyceal afferent terminals within the semi-intact epithelium as described^9^. The EPC-10 (HEKA) patch clamp amplifier was controlled by Patchmaster software (Molecular Devices). In voltage clamp mode, we recorded currents evoked by iterated voltage steps with the amplifier’s 4-pole low-pass Bessel filter corner frequency at 6 kHz and a sampling interval of 5-25 µs. Capacitive currents were nulled on-line with Patchmaster. Series resistances ranged from 4-32 MΩ (mean 10.4 ± 0.6 MΩ, n = 57) and were compensated on-line by 80.1 ± 0.3%. We also recorded voltages evoked by iterated current steps in current clamp mode. Potentials are corrected for a liquid junction potential of 4 mV, calculated with JPCalc software^11^ as implemented by Clampex 10 (Molecular Devices). Cells were held at –64 mV (in voltage clamp) and resting potential (in current clamp) unless otherwise noted. For steady-state analyses of voltage-dependent properties, command potentials were corrected for series resistance errors.

Data were analyzed with OriginPro 2017 (OriginLab, Northampton MA). Results are presented as means ± SE, medians and/or ranges. Comparisons were made with 2-factor ANOVA, with one factor being genotype (control vs. mutant) and the other epithelial zone (striolar vs. extrastriolar), followed by Tukey’s post-hoc test of significance and estimates of statistical power. Effect size was estimated by Cohen’s d statistic.

**Measurement of vestibular evoked potentials (VsEP).** VsEP was measured in *Cyp26b1^lox/+^* (n = 5), *Foxg1^Cre^;Cyp26b1^lox/+^* (n = 10) and *Cyp26b1* cKO (n = 9) mice. VsEP recordings were based on the methods published previously^12-14^. Briefly, animals were anesthetized with a ketamine/xylazine (18:2 mg/ml) mixture, 7 μl per gram body weight, injected intraperitoneally. Core body temperature was maintained at 37.0 ± 0.2°C using a homoeothermic heating pad system. Vestibular stimuli were delivered by securing the mouse head to a mechanical shaker using a noninvasive head clip. Linear acceleration pulses (17 pulses/s, 2 ms duration) ranging from +6 to −18 dB re: 1.0 g/ms (where 1 g = 9.8 m/s^2^), adjusted in 3 dB steps were presented to the head in the naso-occipital axis. Subcutaneous needle electrodes were placed posterior to the right pinna and at the right hip for inverting and ground electrodes, respectively. Stainless-steel wire placed subcutaneously at the nuchal crest served as the noninverting electrode. Electroencephelographic activity was amplified (200,000×), filtered (300–3000 Hz), and digitized (1024 points at 10 μs/point). Two hundred fifty-six primary responses were averaged and replicated for each VsEP waveform. A VsEP intensity series was collected beginning at the maximum stimulus level (i.e., +6 dB re: 1.0 g/ms) with and without acoustic masking (50–50,000 Hz forward masker at 90 dB SPL), and then descending in 3 dB steps until no response was visible. The first positive (P1) and negative (N1) response peaks of the VsEP waveform were scored for each intensity level. Thresholds (measured in dB re: 1.0 g/ms) was obtained from the scored VsEP waveforms.

**Measurement of horizontal angular vestibulo-ocular reflexes (aVOR).** aVOR was measured in *Foxg1^Cre^;Cyp26b1^lox/+^* (n = 5) and *Cyp26b1* cKO (n = 5) mice. Techniques for measurement of aVOR eye movements in response to whole-body head rotations in alert mice were described elsewhere^15^. Briefly, under inhalational and local anesthesia, the dorsal cranium was exposed and dental adhesive was used to affix a head post for animal restraint during subsequent testing performed after recovery from anesthesia. The post was oriented so that when the animal was placed into a restraining device atop a servo-controlled rotating table, the plane tangent to the flat part of the dorsal skull was pitched 30° ‘nose-down’ from Earth horizontal. The axes of the animal's horizontal semicircular canals aligned to within 10° of the Earth-vertical axis.

Eye movements were measured using marker-based 3-dimensional video-oculography. Marker arrays fashioned from photo paper saturated with fluorescent yellow ink were opaque except for three fluorescent 200×200 μm windows separated by 200 μm and arranged in a 45° right triangle. Images were acquired at a rate of 180 frames/s with 500×400 pixel frame size and resolution 263 pixels/mm at the eye surface, equating to resolution of resolution <0.3° for eye angular position. Data were interpolated on a 1 kHz time base using a nonlinear filter based on a running spline (LabVIEW). We adjusted the spline smoothness parameter so that the correlation coefficient R^2^ between the raw and spline-filtered data was greater than 0.80.

For rotational testing, the center of the animal's skull was approximately aligned with the motor's Earth-vertical axis. Transient yaw whole body rotation stimuli were delivered at 3000°/s^2^ constant acceleration for 100 ms to a peak/plateau velocity of 300°/s lasting 400 ms, followed by a 3000°/s^2^ deceleration for 100 ms to rest at 90° from the starting position. Sinusoidal stimuli were 0.02-10 Hz at peak velocity 100°/s for ≥10 cycles per trial. Eye rotation data were converted to rotation vectors in head coordinates and analyzed as described previously^16^. Yaw (horizontal) head angular velocity data were inverted prior to gain calculation. For transient stimuli, we restricted analysis to slow phase nystagmus responses during the 100 ms constant-acceleration time segment after onset of each stimulus. For each trial, response latency was computed as the time difference between the zero-velocity-intercept times for lines fit in a least-mean-square sense to eye and head velocity during the constant-acceleration stimulus portion of the stimulus. Using the slopes of the same fitted lines, we computed “constant acceleration segment gain” G_A_ as the mean over all cycles of the ratio of eye acceleration to head acceleration. For sinusoidal stimuli, we removed quick phases and saccades manually prior to further analysis of slow phase nystagmus data, for which ≥10 cycles per trial were averaged and used to compute gain and phase of the yaw eye angular velocity response relative to the head angular velocity stimulus. Positive phase lead denotes a rightward slow phase nystagmus response leading a leftward head rotation stimulus. Sinusoidal frequency response data were further parameterized by fitting a first order high pass filter to each animal’s data. The resulting gain and corner frequency parameters were used for statistical comparison between mouse groups. Results are expressed as mean ± SEM. Gain and latency data were analyzed using a Mann Whitney U with significance set at P = 0.05.

**Off-vertical axis rotation (OVAR) testing.** *Foxg1^Cre^;Cyp26b1^lox/+^* controls (n = 6) and *Cyp26b1* cKO mice (n = 6), ranging between 6 to 7 months old, were used. Techniques employed for measurement of eye movements during off-vertical axis rotation in alert mice were described elsewhere^17^**.** Briefly, recordings made after fixating mice on a rotating platform, which was tilted 17 degrees with respect to the ground. Platforms speed was increased from 0 to 50 deg/sec in 500 miliseconds and maintained its constant velocity for 72 seconds (10 complete rounds) before being stopped. Eye movements were measured using video occular-graphy (iScan). Eye velocity signal including peak velocity, time constant, amplitude and phase of the sinusoidal component, sinusoidal frequency, and the offset of the final eye velocity was measured in each group.

**Behavioral tests.** In all experiments, *Foxg1^Cre^;Cyp26b1^lox/+^* mice were used as controls. Both males and females were included in behavioral tests and all tests were conducted at NIDCD vestibular core facility, NIMH behavioral core facility and Johns Hopkins University.

*Open field test.* Three-month old control (n = 6) and *Cyp26b1* cKO mice (n = 5) were used. Each mouse was placed in an open arena. Two minutes after habituation, subject was video recorded for 5 min. Trace of each mouse was visualized followed by analysis with Topscan software (Clever Sys Inc.).

*Forced swim test.* Two-month old control (n = 9) and *Cyp26b1* cKO mice (n = 15) were used. Mice were placed in a container of warm water ranging in temperature between 24-30°C in either light or dark condition and their ability to swim was recorded and scored as previously described^18^.

*Rotarod test.* Controls (n = 10) and *Cyp26b1* cKO mice (n = 9), ranging from 2 to 5 months old, were used. Each mouse was placed on a motorized rotating rod (ROTA ROD, Panlab, Harvard Apparatus) that gradually accelerated from 5 to 40 rpm in 5 min, and the time required for the mouse to fall off the rotarod was scored. Each mouse underwent tests for three consecutive days with 5 trials per day. First day was considered to be the training day. Averaged score for each day was processed for statistical analysis.

*Balance beam test on a 20 mm-wide beam.*

Control (n = 5) and *Cyp26b1* cKO mice (n = 6), ranging between 6 to 7 months old, were used. The balance beam apparatus consisted of a 60 cm long and 20 mm-wide beam that was positioned 70 cm above ground with an escape box on one end. Walking speed was measured by recording the time the animal took to reach the escape box from the opposite end of the beam.

*Balance beam test on a 6 mm-wide beam.* Control (n = 5) and *Cyp26b1* cKO mice (n = 6), ranging between 6 to 7 months old, were used. The balance beam apparatus consisted of a 80 cm long and 6 mm-wide beam, situated 70 cm above the ground. Mice were placed at the midpoint of the beam and the time taken to reach the endpoint on either side (time to traverse 40 cm) was measured. Mice were scored ‘’time out’’ when failed to reach the endpoint in 2 minutes.

**Head tremors measurements of P9 pups.** P9 controls (n = 8) and *Cyp26b1* cKO (n = 6) pups were placed in an open arena. Their motor activities and behavior were video-recorded for 10 min. The total distance traversed, the number of bouts per 100 mm traveled and duration of each bout episode were measured using the Topscan software.

**Head tremor measurements of adult mice.** Control (n = 6) and *Cyp26b1* cKO mice (n = 6) between 6 to 7 months old were used. Head movements were recorded using a miniture head motion sensor affixed on the top of the skull, which comprises a 3D accelerometer (measures linear acceleration; right/left, fore/aft, and up/down) and 3D gyroscope (measures angular velocity: pitch, roll, and yaw). Recordings were made while experimental mice were at rest, placed within a cylinder (9 cm diameter and 21.5 cm height) that limited their motion.

**Statistical analysis.**

T-test was used for comparison of two-samples, and either one-way or two-way ANOVA followed by Tukey’s multiple comparison tests was used for more than two samples. Data distribution was assumed to be normal but not formally tested. All statistics were conducted by Prism 7 (GraphPad inc) except in cases described separately. All data are shown as average ± SEM.

**SUPPLEMENTAL LEGENDS**

**Supplemental figure 1. Complementary expression patterns of *Cyp26b1* and *Aldh1a3* in the saccule.**

(**a**-**c**) Whole mount *in situ* hybridization analyses of *Cyp26b1*, *Aldh1a3*, and *β-tectorin* in the saccule (sac) at E18.5. *Cyp26b1*-positive region corresponds to the *β-tectorin*-positive striolar region. *Aldh1a3* is predominantly expressed in the peripheral region. (**d**-**f**) Adjacent sections of E15.5 saccule showing complementary expression pattern of *Cyp26b1* and *Aldh1a3*. Scale bars for both whole mount and section images are 200 μm. A, anterior; L, lateral; D, dorsal.

**Supplemental figure 2. Disruption of RA signaling affects striolar formation in the saccule.**

(**a**-**d**) Immunohistochemistry and quantification of oncomodulin (Ocm)^+^ HCs in E18.5 saccules (sac). Ocm^+^ striolar type I HCs is reduced in the *Cyp26b1^-/-^* saccule (**b**, **d**, 3.1 ± 0.7 %, n = 3, P = 0.0003) but increased in the *Aldh1a3^-/-^* saccule (**c, d,** 34.6 ± 2.3 %, n = 5, P < 0.0001), compared to control saccule (**a**, **d**, 17.7 ± 2.6 % in controls, n = 6)

(**e**-**h**) Immunohistochemistry and quantification of β-tectorin^+^ SCs in E18.5 saccules. β-tectorin^+^ SC area is not changed in either the *Cyp26b1^-/-^* (**f**, **h**, 18.5 ± 0.8 %, n = 3, P = 0.8464) or in the *Aldh1a3^-/-^* saccule (**g**, **h**, 16.2 ± 0.8 %, n = 4, P = 0.6655), compared to control saccule (**e**, **h**, 17.4 ± 1.3 % in controls, n = 4). (**i**-**k**) Immunostaining and quantification of calbindin^+^ afferent neurons in P40 saccules. Calbindin expression detected in striolar region of control saccules (**i**, **k**, 26.6 ± 2.1 % in controls, n = 3) is reduced in *Cyp26b1* cKO saccules (**j**, **k**, 12.1 ± 0.7 % in controls, n = 3, P = 0.0036). The one-way ANOVA with multiple comparisons was applied. ***P* < 0.01 and ****P* < 0.001. Scales bar; 200 μm. D, dorsal; A, anterior.

**Supplemental figure 3. Differential response to loss of *Aldh1a3* in the utricles and saccules.**

(**a-f**) Immunohistochemistry and quantification of Ocm^+^ HCs in E18.5 utricles (ut) and saccules (sac). (**a**-**c**) Ocm, which labels striolar type I HCs in the control utricles, is only expressed medial (M-LPR, 30.2 ± 2.8%, n = 4) but not lateral (L-LPR, 1.9 ± 0.9%) to the line of polarity reversal (LPR) (**a**, **c**). Ocm staining in *Aldh1a3*^-/-^ utricles is increased in the medial (47.3 ± 2.8%, n = 3, P = 0.0048) but not the lateral region (0.6 ± 0.2%, P = 0.7101) (**b**, **c**). (**d-f**) In control saccule, striola straddles the LPR with Ocm^+^ HCs being expressed in inner (IR, 20.1 ± 0.6%, n = 3) and outer (OR, 25.2 ± 0.7%) regions of saccules (**d**, **f**). In contrast to the utricle, Ocm-positive region is expanded on both IR (33.4 ± 0.7%, n = 3, P = 0.0011) and OR (40.4 ± 0.5%, P = 0.0001) of the LPR in *Aldh1a3^-/-^* saccules (**e**, **f**). Unpaired t-test was applied. ***P* < 0.01 and ****P* < 0.001. Scale bar; 200 μm. A, anterior; M, medial; D, dorsal.

**Supplemental figure 4. Loss of regional difference in hair cell density in *Cyp26b1* cKO utricles.**

(**a**-**c**) Immunohistochemistry and quantification of Myosin7a^+^ (magenta) HCs in P0 utricles. Ocm (green) marks control striola. The posterior half of the middle-third region of each utricle is sub-divided into four regions. In area 1 of control utricles, which represents the lateral extrastriola (LES), HCs are smaller and densely packed (**a’**, **c**, 56.5 ± 3.5 per 0.01 mm^2^, n = 4). HCs are larger and wider apart in area 2, which represents striolar region in controls (**a’’**, **c**, 41.7 ± 1.9 per 0.01 mm^2^, P = 0.0057). In *Cyp26b1* cKO mutant utricles, area 2 have smaller and denser HCs (**b’’**, **c**, 63.2 ± 6.2 per 0.01 mm^2^, n = 4,), similar to HCs in area1 (**b’**, **c**, 60.0 ± 4.6 per 0.01 mm^2^, P = 0.3457). (**d**) Averaged ratio of HC density in area 2 to area 1 in control (0.74 ± 0.03), is smaller than *Cyp26b1* cKO mutant utricles (1.05 ± 0.07, P = 0.0037). a, anterior; m, medial. The two-way ANOVA with multiple comparisons was applied to (**c**). Unpaired t-test was applied to (**d**). A, anterior; M, medial. ***P* < 0.01. Scale bar; 200 μm for (**a**), and 30 μm (**a’’**).

**Supplemental figure 5. Position of the line of polarity reversal (LPR) is maintained in *Cyp26b1^-/-^* utricles and saccules.**

(**a**-**f**) Immunohistochemistry and quantification of area of Spectrin (magenta), which labels the cuticular plate of HCs and identifies the hair bundle orientation^19^, in E18.5 utricles and saccules of controls and *Cyp26b1^-/-^* ears. Ocm (green) labels control striola. In control utricles, the LPR (white line in **a**) is located lateral to the Ocm^+^ striola, whereas it bisects the striola in the saccule (white line in **c**). The LPR approximately halves area of utricles (61.6 ± 2.1% M-LPR, n = 3) and saccules (52.6 ± 1.3% IR, n = 3). In *Cyp26b1* mutants, the position of the LPR remains in its relative position in both utricle (57.1± 1.6% M-LPR, n = 3, P = 0.1609) and saccule (52.8 ± 2.6% IR, n = 3, P = 0.9600) despite the loss of Ocm immunostaining. A, anterior; M, medial; D, dorsal. Unpaired t-test was applied. Scale bar, 200 μm.

**Supplemental figure 6. Loss of regional difference in K/S ratio of hair bundles in *Cyp26b1* cKO utricles.**

(**a**-**c**) Immunohistochemistry and measurements of the ratio of lengths of the kinocilium to tallest stereocilia (K/S ratio) in P10 utricles of control and *Cyp26b1* cKO mutant ears. The utricles were labeled with phalloidin (magenta, labeling stereocilia) and antibodies to Arl13b (green, specific for labeling the kinocilium). The LPR was used to approximate the putative striolar region. In the striola of control utricles (**a’**, horizontal flip of the inset in **a**), the height of the kinocilium and the tallest stereocilium of a hair bundle is similar to each other (**a’’**, **c**, K/S ratio: 0.98 ± 0.03, n = 12 hair bundles), whereas the kinocilium is longer than the tallest stereocilium in HCs of L-LPR (**a’’’**, **c**, K/S ratio: 1.24 ± 0.04, n = 16, P = 0.0010). In contrast, kinocilium lengths of HCs in the striolar region of mutant utricles (**b’**) are longer than the tallest stereocilium (**b’’**, **c**, K/S ratio: 1.29 ± 0.04, n = 14), which are comparable to those in L-LPR (**a’’’**, **b’’’**, P = 0.6438). Green and magenta arrowheads indicate the approximate tips of kinocilium and the tallest stereocilium, respectively. The one-way ANOVA with multiple comparisons was applied. ***P* < 0.01. L, lateral; A, anterior. Scale bar; 200 μm for **a**, 30 μm for **a’**, and 10 μm **a’’**.

**Supplemental figure 7. No hyperactivity in *Cyp26b1* cKO mice.** (**a**) Quantification of open field tests for control and mutant mice. Measurements of total distance a mouse traveled over a 5-minute period when placed in the corner of an open field box. No difference between controls and mutants was observed.

**Supplementary video 1**. **Open field behavior of *Cyp26b1*** **cKO mice.**

No obvious behavioral problems except slight head tremors were detectable in mutants.

**Supplementary video 2. Adult mice traversing on a balance beam.**

Compared to controls, *Cyp26b1* cKO mice exhibit difficulty in traversing on a 6 mm narrow beam.

**Supplementary video 3. Head tremors of *Cyp26b1* cKO mice at P9.**

*Cyp26b1* cKO mice exhibit increased head tremors during self-motion.

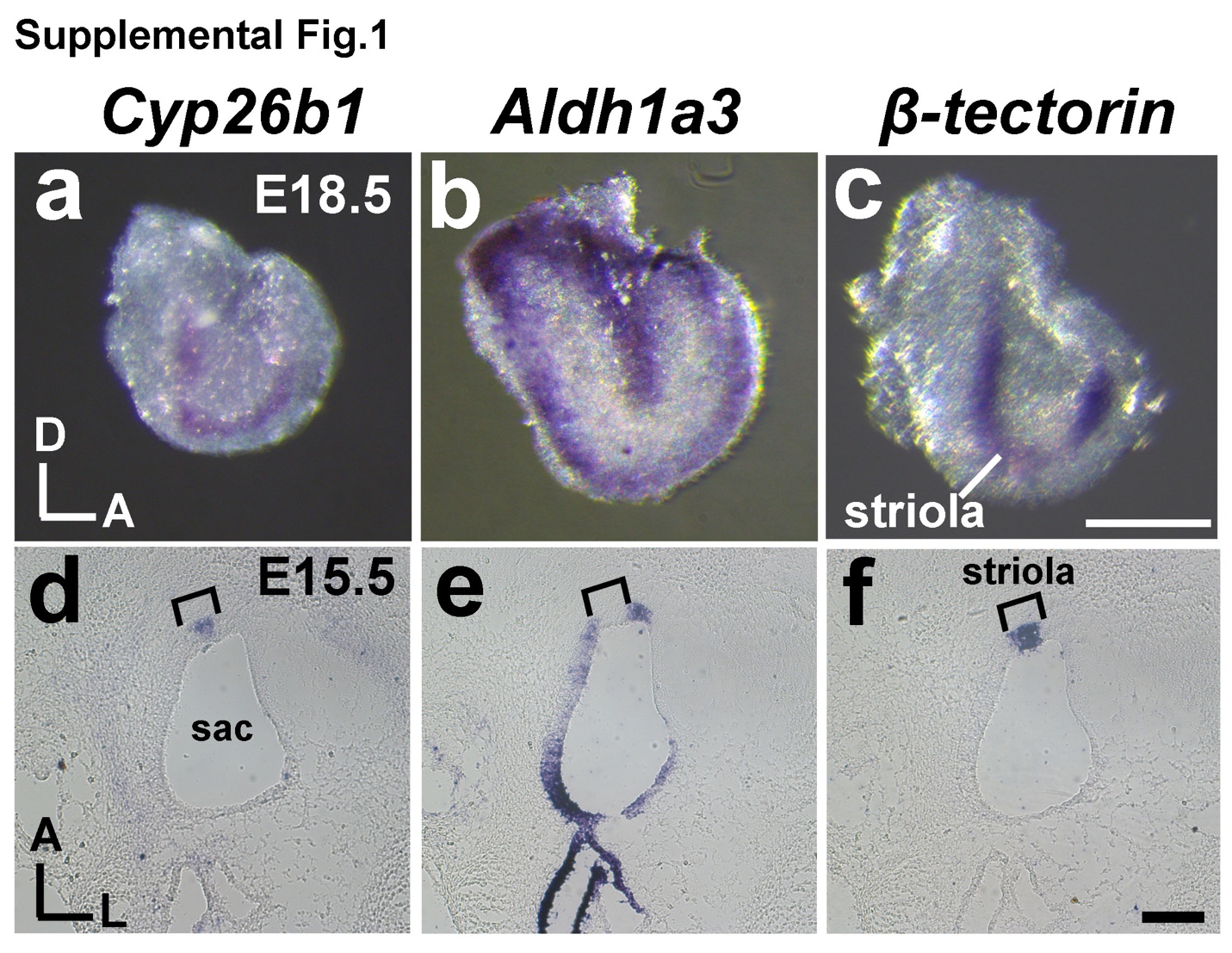

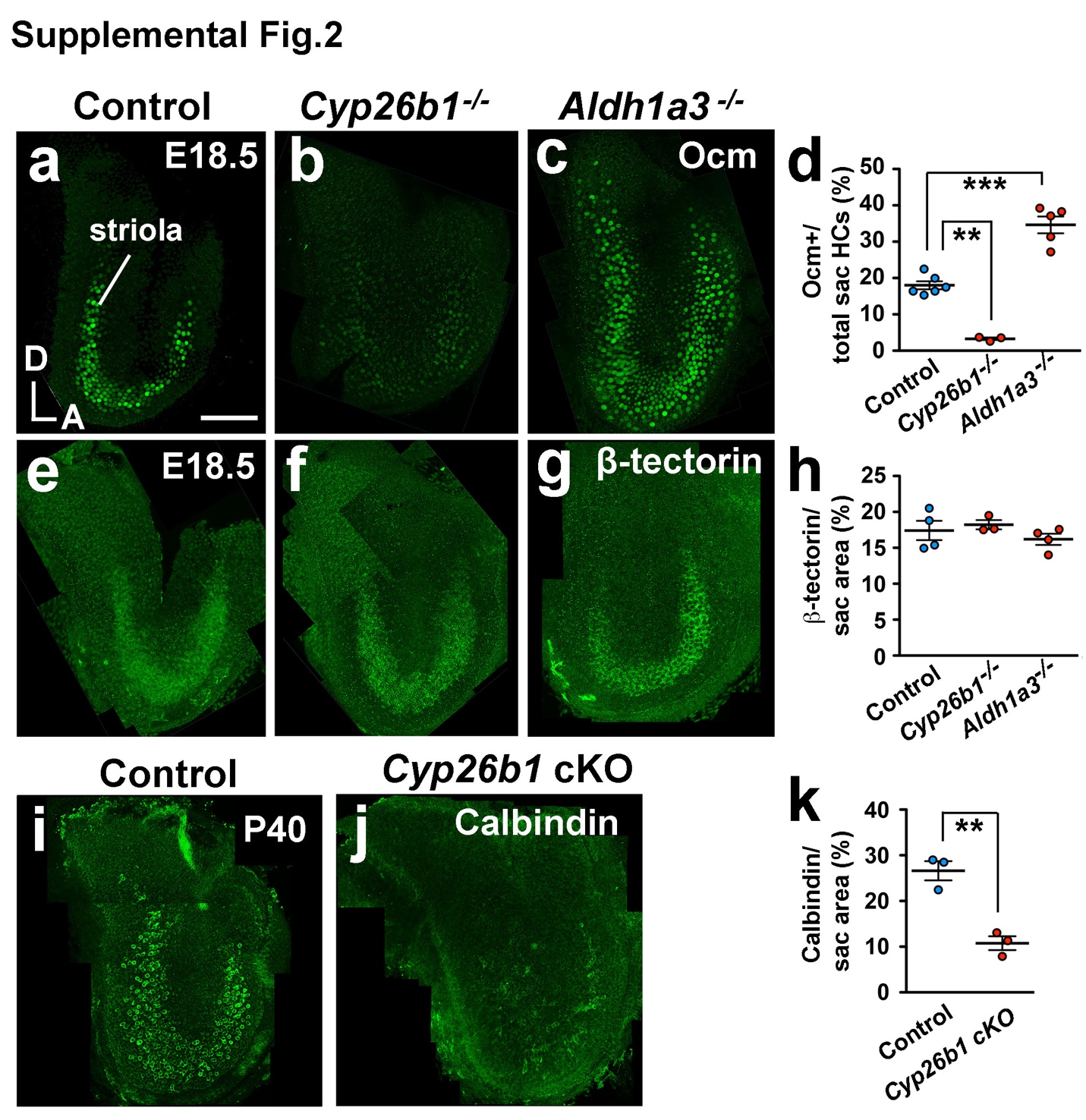

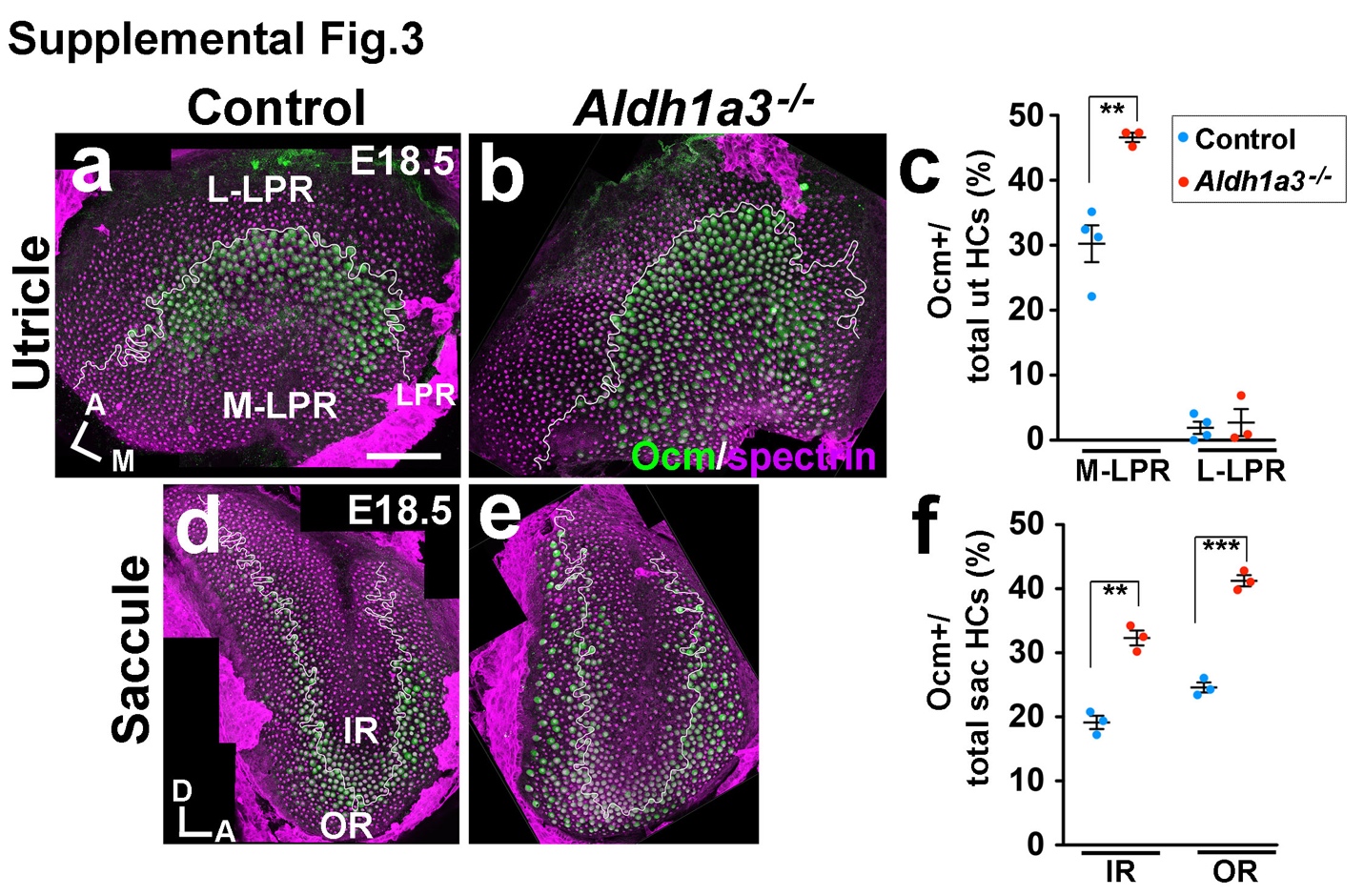

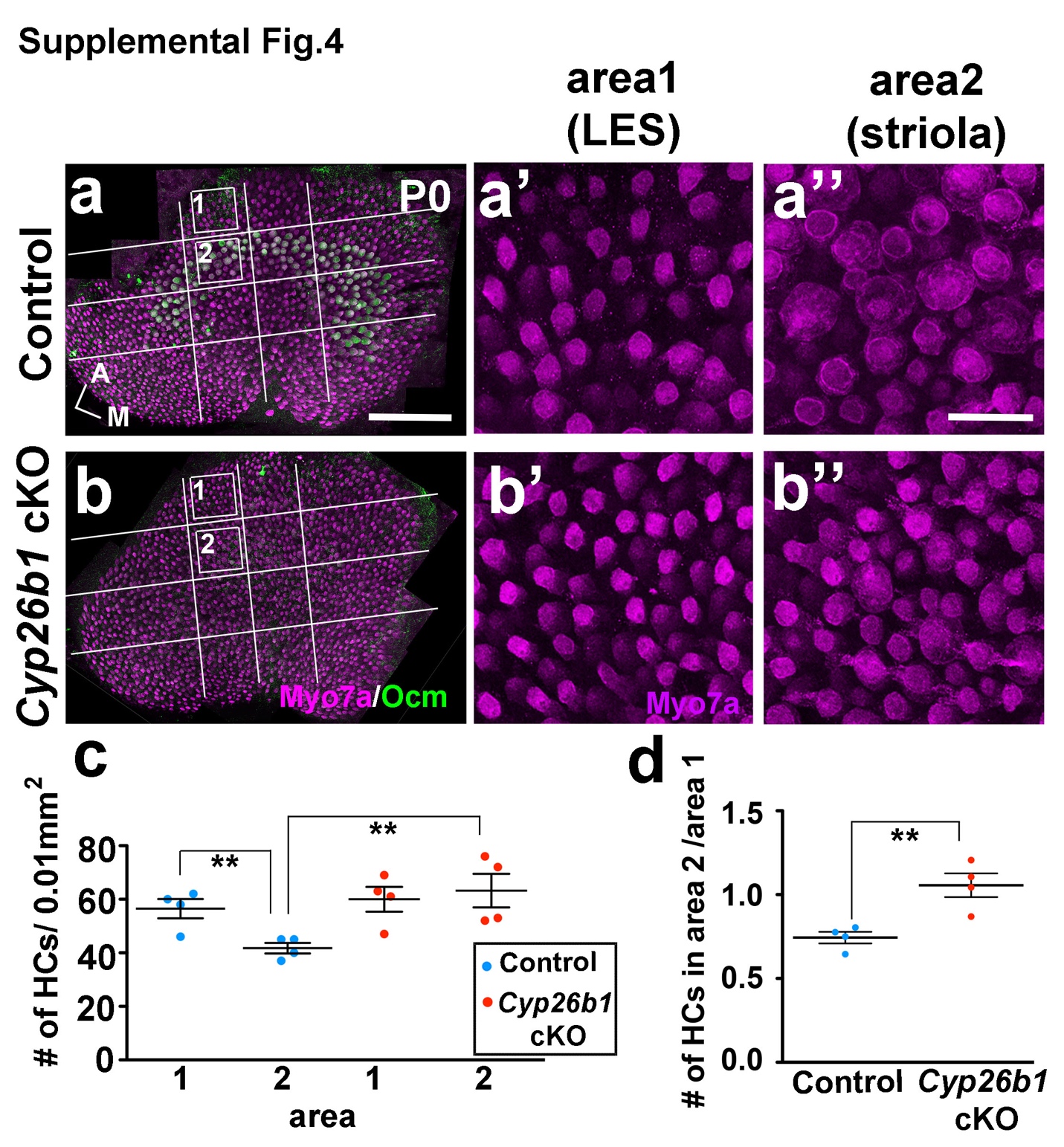

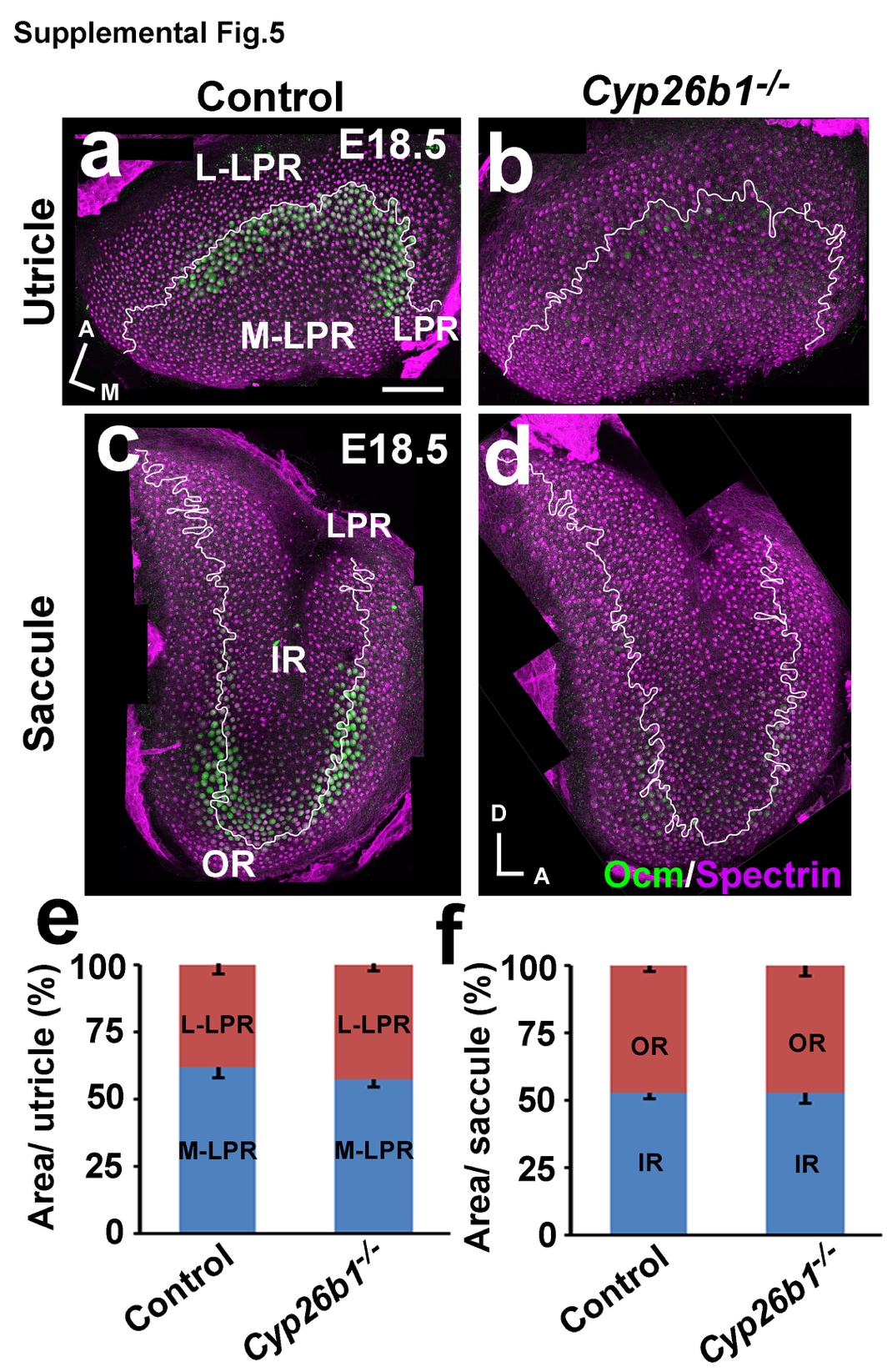

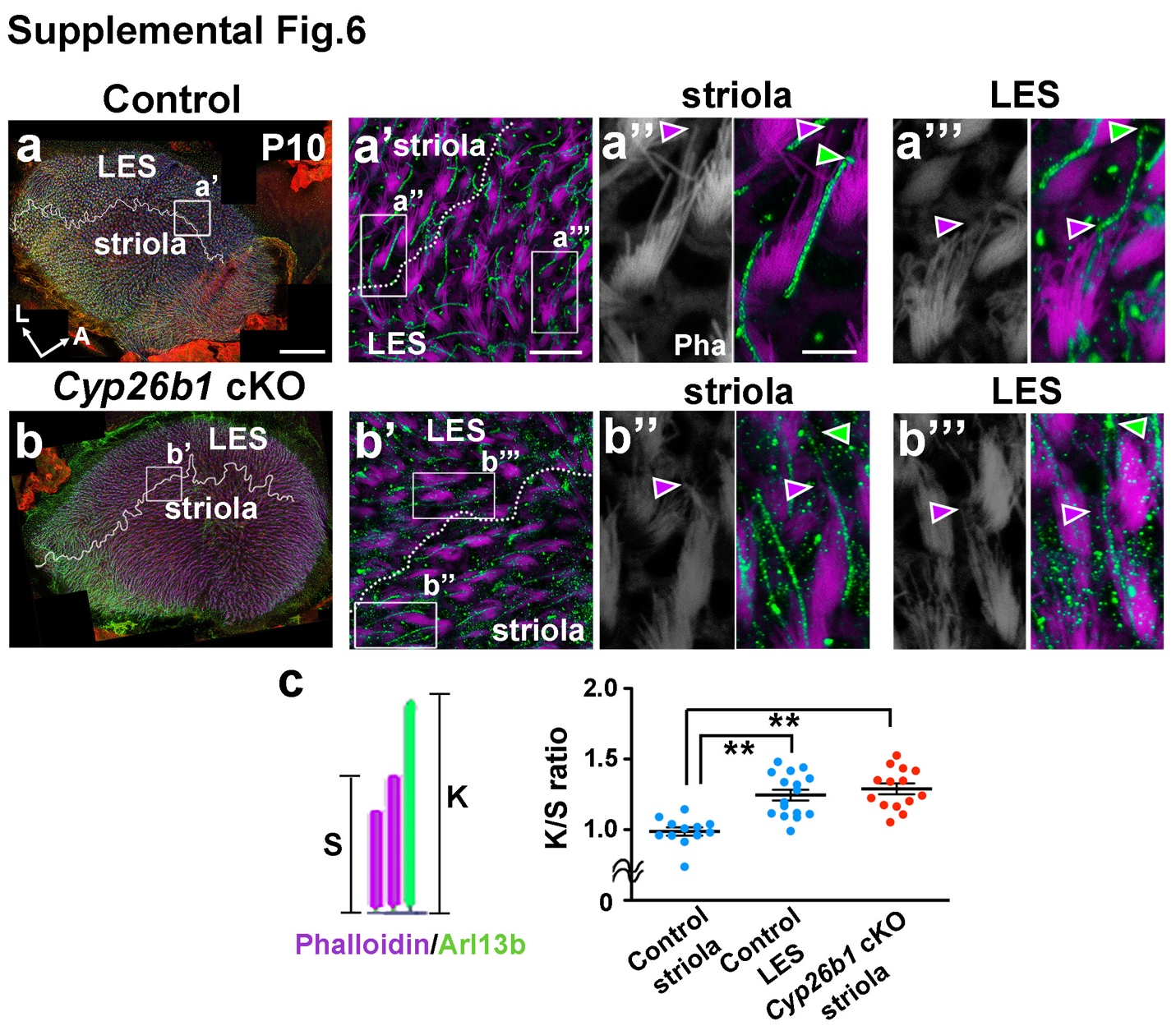

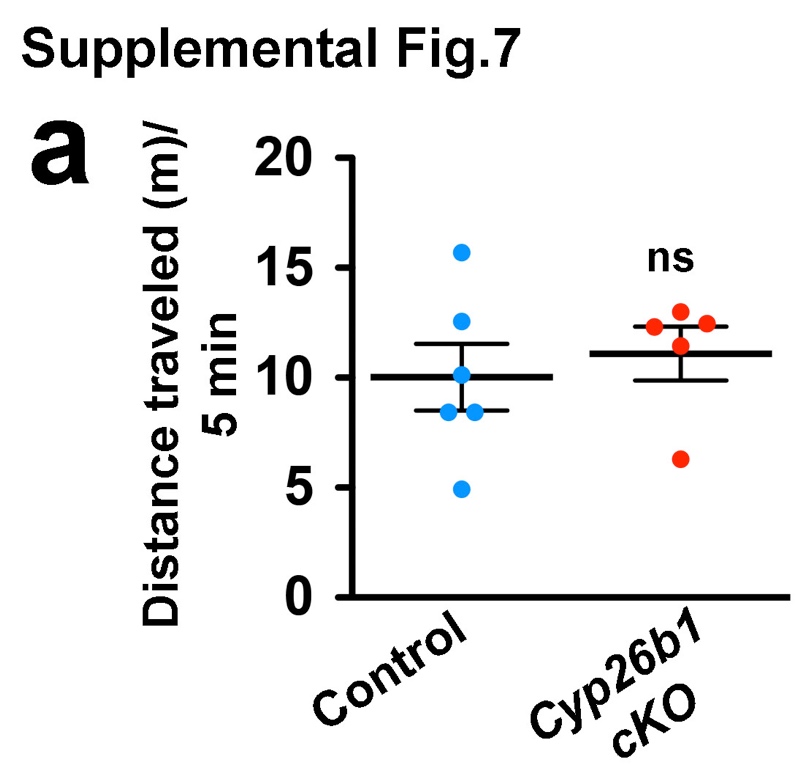

References

1	Molotkov, A., Molotkova, N. & Duester, G. Retinoic acid guides eye morphogenetic movements via paracrine signaling but is unnecessary for retinal dorsoventral patterning. *Development* **133**, 1901-1910, doi:10.1242/dev.02328 (2006).

2	Hebert, J. M. & McConnell, S. K. Targeting of cre to the Foxg1 (BF-1) locus mediates loxP recombination in the telencephalon and other developing head structures. *Dev Biol* **222**, 296-306, doi:10.1006/dbio.2000.9732 (2000).

3	Morsli, H., Choo, D., Ryan, A., Johnson, R. & Wu, D. K. Development of the Mouse Inner Ear and Origin of Its Sensory Organs. *J. Neurosci.* **18**, 3327-3335 (1998).

4	Rau, A., Legan, P. K. & Richardson, G. P. Tectorin mRNA Expression Is Spatially and Temporally Restricted During Mouse Inner Ear Development. *The Journal of Comparative Neurology* **405**, 271-280 (1999).

5	Zhao, X. *et al.* Retinoic acid promotes limb induction through effects on body axis extension but is unnecessary for limb patterning. *Curr Biol* **19**, 1050-1057, doi:10.1016/j.cub.2009.04.059 (2009).

6	Forge, A., Taylor, R. R., Dawson, S. J., Lovett, M. & Jagger, D. J. Disruption of SorCS2 reveals differences in the regulation of stereociliary bundle formation between hair cell types in the inner ear. *PLoS Genet* **13**, e1006692, doi:10.1371/journal.pgen.1006692 (2017).

7	Dupe, V. *et al.* A newborn lethal defect due to inactivation of retinaldegyde dehydrogenase type 3 is prevented by maternal retinoic acid treatment. *PNAS* **100**, 14036-14041 (2003).

8	Jiang, T., Kindt, K. & Wu, D. K. Transcription factor Emx2 controls stereociliary bundle orientation of sensory hair cells. *Elife* **6**, doi:10.7554/eLife.23661 (2017).

9	Songer, J. E. & Eatock, R. A. Tuning and timing in mammalian type I hair cells and calyceal synapses. *J Neurosci* **33**, 3706-3724, doi:10.1523/JNEUROSCI.4067-12.2013 (2013).

10	Rusch, A. & Eatock, R. A. A delayed rectifier conductance in type I hair cells of the mouse utricle. *J Neurophysiol* **76**, 995-1004 (1996).

11	Barry, P. H. JPCalc, a software package for calculating liquid junction potential corrections in patch-clamp, intracellular, epithelial and bilayer measurements and for correcting junction potential measurements. *J of Neuroscience Methods* **51**, 107-116 (1994).

12	Vijayakumar, S. *et al.* Vestibular dysfunction, altered macular structure and trait localization in A/J inbred mice. *Mamm Genome* **26**, 154-172, doi:10.1007/s00335-015-9556-0 (2015).

13	Vijayakumar, S. *et al.* Rescue of peripheral vestibular function in Usher syndrome mice using a splice-switching antisense oligonucleotide. *Hum Mol Genet* **26**, 3482-3494, doi:10.1093/hmg/ddx234 (2017).

14	Mock, B., Jones, T. A. & Jones, S. M. Gravity Receptor Aging in the CBA/CaJ Strain: A Comparioson to Auditory Aging. *JARO* **12** (2011).

15	Webb, S. W. *et al.* Regulation of PCDH15 function in mechanosensory hair cells by alternative splicing of the cytoplasmic domain. *Development* **138**, 1607-1617, doi:10.1242/dev.060061 (2011).

16	Migliaccio, A. A., Macdougall, H. G., Minor, L. B. & Della Santina, C. C. Inexpensive system for real-time 3-dimensional video-oculography using a fluorescent marker array. *J Neurosci Methods* **143**, 141-150, doi:10.1016/j.jneumeth.2004.09.024 (2005).

17	Beraneck, M., Bojados, M., Le Seac'h, A., Jamon, M. & Vidal, P. P. Ontogeny of mouse vestibulo-ocular reflex following genetic or environmental alteration of gravity sensing. *PLoS One* **7**, e40414, doi:10.1371/journal.pone.0040414 (2012).

18	Hardisty-Hughes, R. E., Parker, A. & Brown, S. D. A hearing and vestibular phenotyping pipeline to identify mouse mutants with hearing impairment. *Nat Protoc* **5**, 177-190, doi:10.1038/nprot.2009.204 (2010).

19	Deans, M. R. *et al.* Asymmetric distribution of prickle-like 2 reveals an early underlying polarization of vestibular sensory epithelia in the inner ear. *J Neurosci* **27**, 3139-3147, doi:10.1523/JNEUROSCI.5151-06.2007 (2007).

20	Li, A., Xue, J. & Peterson, E. H. Architecture of the mouse utricle: macular organization and hair bundle heights. *J Neurophysiol* **99**, 718-733, doi:10.1152/jn.00831.2007 (2008).
